# Deaf1 regulates the coordinated repression of synaptic genes to limit synaptogenesis

**DOI:** 10.1101/2024.10.30.621128

**Authors:** James A. Kentro, Gunjan Singh, Tuan M. Pham, Justin Currie, Saniya Khullar, Audrey T. Medeiros, Erica Larschan, Kate M. O’Connor-Giles

## Abstract

Synapses are the primary sites of communication in the nervous system. Synapse formation requires the simultaneous expression of hundreds of proteins across cells. Through genomic and functional studies in *Drosophila*, we investigated the regulatory mechanisms underlying the spatiotemporal coordination of synaptic gene expression during neurodevelopment. We find that chromatin opening is coordinated across hundreds of synaptic genes immediately preceding transcription, whereas repression of these genes outside developmental periods of synaptogenesis is regulated by distinct mechanisms. We identify the conserved DEAF1 transcription factor (TF) as a direct repressor of synaptic genes. Disruption of DEAF1 or pioneer TF CLAMP, which promotes DEAF1 binding, increases synaptic gene expression across neuronal subtypes, leading to excess synapse formation. DEAF1, which is linked to syndromic intellectual disability, is also sufficient to block synaptogenesis. These findings reveal the importance of coordinated synaptic gene repression in sculpting neuronal connectivity and broaden our understanding of gene regulation in neurodevelopment.

## Introduction

The nervous system is remarkable for its cellular diversity and functional complexity. In the human brain, billions of neurons of diverse subtypes form trillions of synapses that connect neurons into the functional circuits that interpret the environment and control behavior. There has been significant progress in understanding the role of transcriptional regulators in generating the great diversity of neurons(*1–4*). Yet, how gene regulatory networks (GRNs) direct the development of shared neuronal traits has been far less studied.

The chemical synapses that underlie neuronal communication are hallmarks of mature neurons. Synapses are complex intercellular structures designed for the rapid transmission of information in the form of Ca^2+^-dependent neurotransmitter release and uptake. Building a functional synapse necessitates the simultaneous expression of hundreds of proteins, including neurotransmitter transporters and receptors, regulators of synaptic vesicle release, and scaffolding and adhesion molecules, as well as the proteins that regulate their function and transport to synapses(*5*, *6*). Adding to the challenge, the expression of synaptic genes, which we define broadly here as all genes that encode proteins required for the formation of functional synapses, must not only be coordinated within a given cell, but across synaptically connected neurons. While this coordination likely occurs at multiple levels, our analysis of *Drosophila* time-series transcriptomic data indicates that ∼500 synaptic genes encoded at hundreds of different genomic loci are temporally coordinated at the level of transcription with high levels of expression restricted to developmental windows of synaptogenesis. These synaptic genes include both pan-neuronal and diverse neuron subtype-specific synaptic genes indicating coordination across neuronal subtypes.This broad spatio-temporal coordination is consistent with a recent investigation of transcriptional regulation of circuit assembly in the *Drosophila* visual system, which observed temporal coordination of synaptic genes across neuronal cell types(*7*). Yet, how the temporal coordination of synaptic gene transcription across neuronal subtypes is achieved remains poorly understood.

Here, we investigate the mechanisms and regulators that temporally coordinate the transcription of synaptic genes during neurodevelopment. Consistent with prior studies implicating chromatin regulation in neuronal maturation(*8–11*), we find that chromatin opening is coordinately regulated across hundreds of synaptic genes immediately preceding the onset of synaptic gene expression. In contrast, the coordinated repression of synaptic gene expression observed following peak synaptogenesis is unlinked to chromatin accessibility, prompting us to search for repressive factors. Repressive transcriptional mechanisms are critical for many gene regulatory processes that drive key developmental decisions(*12*, *13*) but remain less understood in the nervous system than activating mechanisms. Activation of neuronal subtype-specific gene expression is mediated by well-defined DNA binding transcription factors (TFs) called terminal selectors(*4*, *14–16*). While pan-neuronal gene expression is less studied, evidence from *C.elegans* indicates that pan-neuronal genes are activated by subtype-specific combinations of TFs acting in concert with broadly expressed TFs(*17*, *18*). In contrast, the molecular mechanisms by which pan-neuronal and subtype-specific synaptic genes are temporally coordinated remains unknown.

Therefore, we combined chromatin accessibility and motif analyses with machine learning approaches to predict GRNs from single-cell multiomic data. We identified DEAF1, which is linked to intellectual disability in humans(*19–21*), and CLAMP, a pioneer factor that regulates chromatin accessibility and promotes polymerase pausing(*22*, *23*), as predicted negative regulators of temporally coordinated synaptic gene expression. DEAF1 binding sites exhibit enriched accessibility at synaptic genes. By measuring the chromatin occupancy and role in regulating gene expression, we find that following peak synaptogenesis DEAF1 directly represses a majority of synaptic genes that are bound and regulated by it. Furthermore, DEAF1 requires CLAMP for its occupancy at chromatin, suggesting a functional relationship. Notably, knockdown of *DEAF1* or *CLAMP* results in excess synapse formation. Conversely, overexpression of *DEAF1* blocks synapse formation, demonstrating that it is necessary and sufficient for repressing synapse formation. These findings support the model that synaptic gene expression is temporally coordinated through the combined action of coordinated chromatin opening across neuronal subtypes, previously identified neuron-specific transcriptional activators(*1–4*), and these newly identified pan-neuronal repressors. Overall, we demonstrate a critical requirement for transcriptional repression in the broad co-regulation of synaptic genes across the developing nervous system and define the importance of these repressive mechanisms in sculpting synaptic connections.

## Results

### Dynamic coordination of synaptic gene transcription across the brain

Coordinated expression of the hundreds of proteins required to form functional synapses could occur at multiple levels, including transcription, translation, and protein trafficking. We identified a number of known synaptic genes with two peaks of transcription correlating with bursts of synaptogenesis during embryonic and adult nervous system development and significant repression of synaptic gene expression outside these windows (Figure 1A), suggesting synaptic genes may be transcriptionally co-regulated. We used the developmental transcriptional expression profiles of 20 well-characterized *Drosophila* genes that encode an array of proteins critical for synapse formation and function (Figure 1A) to generate a ‘synaptic’ transcriptional footprint and compared it to the developmental expression profiles of all *Drosophila* genes(*9*). We identified 485 genes with Spearman correlation scores above 0.8 (Figure 1B, blue; Table S1), all of which are predominantly expressed in the nervous system and enriched for synaptic terms in Gene Ontology (GO) analysis (Figure 1C, dark blue), consistent with the broad coordinated transcription of synaptic genes.

**Figure 1.**
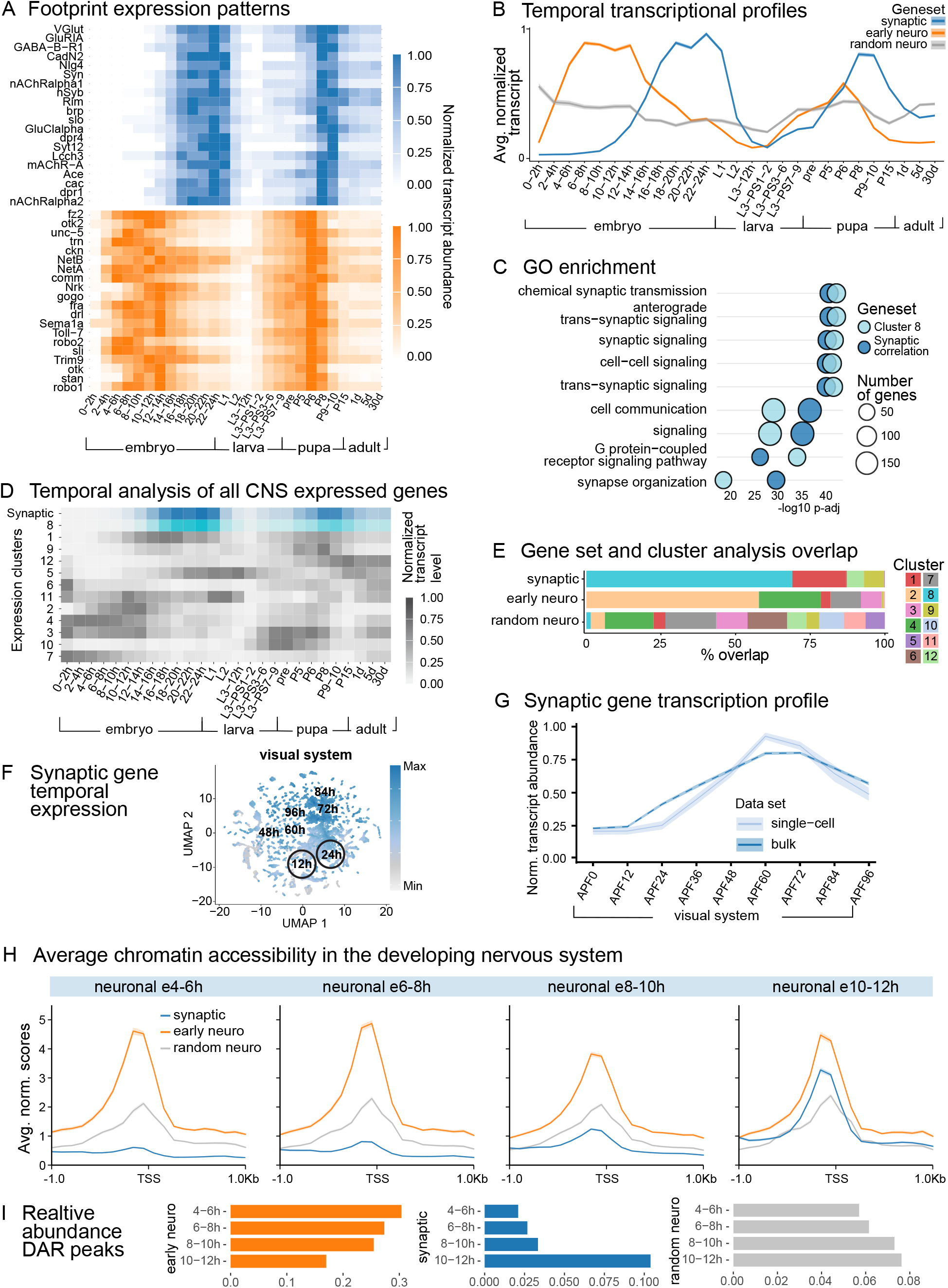
Temporal coordination of synaptic gene transcription and chromatin accessibility. (A) Temporal expression patterns of genes with well characterized synaptic (blue) or neurite development (orange) functions, normalized to peak expression(*9*). (B) Average temporal expression profile of the 485 genes most highly correlated with the footprint patterns in A for synaptic function (blue), early neuronal development (orange), or random nervous system genes (grey). (C) GO analysis of the 485 genes highly coordinated with the synaptic footprint shows enrichment for synapse-related terms; top GO terms for cluster 8 in (D,E). (D) Unbiased k-means clustering analysis of all nervous system expressed Drosophila genes by temporal expression pattern yields one cluster with a pattern closely matching the synaptic pattern (cluster 8, aqua). (E) Gene set/cluster distribution of the 485 synaptic genes mostly map to cluster 8. (F) UMAP of developing pupal visual system scRNA-seq showing composite synaptic gene expression and hours APF(*29*). (G) Temporal pattern of synaptic gene average expression from bulk whole pupa(*9*) and single-cell visual system(*29*) data. (H) Average chromatin accessibility surrounding TSSs of synaptic (blue), neuronal development (orange), and random nervous system genes (grey) in tissue-specific embryonic DNase-seq time-series data(*31*). (I) Abundance of accessible peaks within 3kb of synaptic (blue), early neuronal development (orange), and random nervous system genes (grey) relative to total neuronal peaks at each timepoint. Shading represents ± SEM.

In contrast, genes with lower correlation scores include few well-characterized synaptic genes and are enriched for diverse GO terms (Figure S1A). Most of the 485 genes identified encode proteins with well-characterized roles in synapse formation and/or function, including the core proteins that promote Ca^2+^-dependent synaptic vesicle release at all synapses and the subtype-specific synaptic proteins that regulate the synthesis, transport, and reception of glutamate, acetylcholine, gamma-aminobutyric acid (GABA), and monoamines. 168 of the co-regulated genes remain less characterized in *Drosophila* and are designated as computed genes (CGs). Many encode homologs of proteins linked to synapses in other species and recent studies in *Drosophila* have identified roles in synapse formation and/or function for others, including TRMT9B/Fid(*10*, *11*) and PDZD8(*24*). This temporal pattern appears specific to synaptic genes as a similar analysis of genes associated with neurite development/axon guidance revealed a temporally distinct pattern with peaks of expression that precede peaks of synaptic gene expression (See Figure 1A-B, orange, Table S2), and identified genes enriched for neurite and other early neuronal development GO terms (Figure S1B), consistent with the ordered expression of broad programs of functionally related genes across nervous system development observed across species(*7*). Finally, we observe a similar pattern of synaptic gene expression in pseudobulk embryonic and larval nervous system time-series single-cell RNA-seq data (Figure S1C,D)(*25–27*).

Using an independent and unbiased approach, we performed k-means clustering analysis of all genes expressed in the nervous system based on temporal expression. We identified 12 clusters with distinct temporal expression patterns, one of which closely matches the synaptic gene temporal expression profile defined above (Figure 1D, cluster 8, light blue), and overlaps with 324/485 coordinately regulated synaptic genes (Figure 1E; padj = 0x10^00^). Cluster 8 genes are also similarly enriched for synaptic GO terms (See Figure 1C, light blue). Together, these findings indicate the existence of regulatory mechanisms that temporally coordinate both the transcriptional activation and repression of pan-neuronal and subtype-specific synaptic genes across the nervous system.

We next considered the possibility that the observed dynamic coordination of synaptic gene expression is due to the unique need to build the nervous system twice in pupating animals. To address this, we turned to the *Drosophila* visual system, which develops almost entirely during the pupal stage(*28*). A previous study of pupal time series single-cell RNA-seq in the visual system identified over 180 pan-neuronal genes that were coordinately up-and downregulated across cell types (Pearson’s Correlation Coefficient >.75)(*29*). Notably, these genes were enriched for those encoding synaptic proteins. Using these data, we examined the 485 synaptic genes in our synaptic gene dataset and observed broadly coordinated dynamic coregulation (Figure 1F, G). In the visual system, synaptogenesis initiates around 45h after puparium formation (APF)(*30*). Consistently, synaptic gene transcription initiates by 36h APF with a peak around 60h APF followed by a temporally coordinated decline. The broad repression of synaptic genes following synaptogenesis in the visual system is similar to our observations in the adult CNS (see Figure 1A, B, blue) and suggests that dynamic coordinated activation and repression of synaptic genes may be a general feature of nervous system development.

### Chromatin accessibility changes precede activation of synaptic gene activation

To understand how the expression of hundreds of synaptic genes, which are widely dispersed across chromosomes, is temporally coordinated, we analyzed existing DNase-seq time-series data from isolated neuronal and mesodermal embryonic tissue(*31*) to determine if chromatin accessibility is similarly dynamically regulated at synaptic genes. In early developmental stages, promoter regions upstream of synaptic genes are closed relative to a random set of 485 nervous system expressed genes (Figure 1H, blue vs. grey). Chromatin becomes substantially more accessible at synaptic genes relative to random neuronal genes beginning at 10-12 hours, immediately preceding the coordinated upregulation of synaptic gene transcription. Using time-series proteomic data available for a small subset of synaptic proteins(*32*), we also observe a coordinated increase in synaptic protein abundance immediately following chromatin opening and transcription initiation, suggesting that the coordinated increase in chromatin accessibility ultimately influences protein production (Figure S1E). This dynamic pattern of chromatin accessibility is specific to the nervous system (Figure S1F) and synaptic vs. early neuronal development or random nervous system genes (Figure 1H, blue vs. orange). Furthermore, we determined that synaptic genes have a larger increase in the relative abundance of differentially accessible regions (DARs) at 10-12 hours compared to early neuronal development and random nervous system genes (Figure 1I). It has previously been shown that distinct combinations of TFs can activate pan-neuronal genes in different neurons(*17*, *18*). This finding suggests that broadly coordinated chromatin opening provides a permissive signal that temporally coordinates the action of these factors across the brain during nervous system development.

### Repression of synaptic genes is not driven by changes in chromatin accessibility

To determine if the dynamic regulation of chromatin also corresponds to synaptic gene expression following peak synaptogenesis when transcription is repressed, we investigated chromatin accessibility at these newly accessible sites in later stages of development using larval, pupal, and adult neural ATAC-seq data(*33*, *34*). Strikingly, we found that chromatin remains broadly accessible across synaptic genes throughout these stages (Figure 2A). Nearly all (95%) DARs identified near synaptic genes remain accessible in third instar larval brains when synaptic gene transcription is low. Therefore, repression of synaptic gene expression following peak synaptogenesis is achieved through a distinct mechanism independent of chromatin accessibility changes, possibly to enable rapid activity-dependent re-expression of synaptic genes outside of nervous system development.

**Figure 2.**
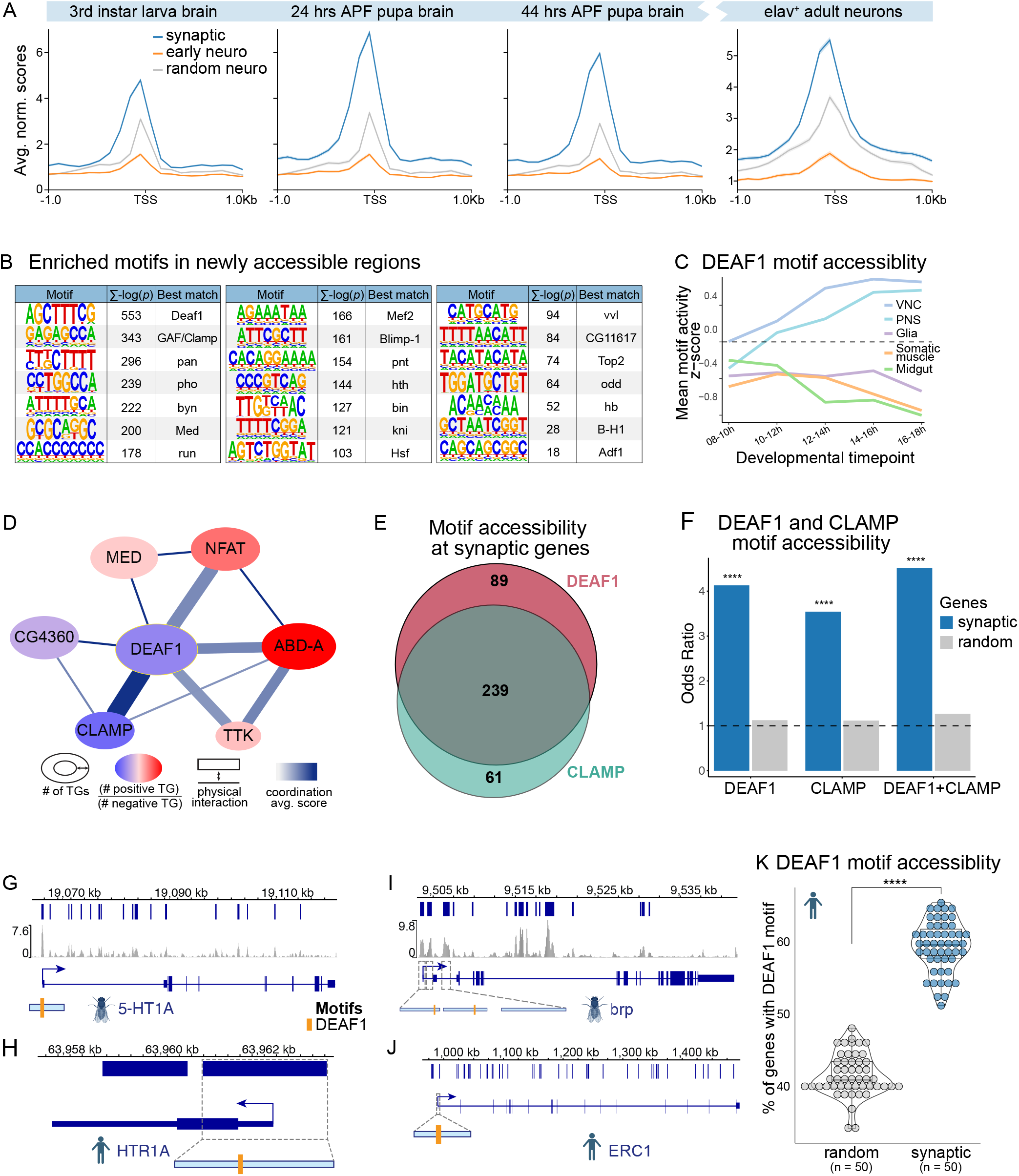
DEAF1 and CLAMP are predicted to co-repress synaptic gene expression. (A) Average chromatin accessibility surrounding TSSs of synaptic genes (blue), neuronal development genes (orange), and random CNS genes (grey) in larval, pupal, and adult ATAC-seq data(33, 34). (B) Enriched motifs in newly accessible regions within 3kb of synaptic genes using Homer(*80*). (C) DEAF1 motif activity scores across cell-classes in embryonic scATAC time-series data(*25*). (D) TF coordination network centered on DEAF1, showing number of predicted synaptic target genes (ellipse size), predicted positive (red) to negative (dark blue) effect ratio on target genes, experimental physical interaction scores (line width), and calculated average weighted correlation score (line color intensity). (E) Overlap of synaptic genes with accessible DEAF1 and CLAMP binding motifs in third instar larval brain ATAC-seq peaks(*33*). (F) Odds ratios of accessible DEAF1 and CLAMP binding motifs at synaptic genes (blue) are enriched beyond random nervous system genes (grey) in larval brain ATAC-seq peaks(*33*). (G-H) DEAF1 motif locations in accessible peaks near TSS of example synaptic genes 5-hydroxytryptamine receptor 1A (5-HT1A, G) and its human ortholog 5-Hydroxytryptamine Receptor 1A (HTR1A, H). (I-J) DEAF1 motif locations in accessible peaks near TSS of the other example synaptic genes bruchpilot (brp, I) and its human ortholog ELKS/RAB6-Interacting/CAST Family Member 1 (ERC1, J). (K) Percent of genes with at least one accessible DEAF1 motif in human cerebral organoid ATAC-seq(*48*) peaks within 3kb of human genes with > 0.8 Spearman correlation to the average temporal single-cell(*47*) expression pattern of orthologs to synaptic footprint genes from Figure 1A. Each point in I represents one timepoint-replicate ATAC-seq sample. In F, ****P<10^-5^ (Fisher’s exact test, Bonferroni p-adjustment). In I, ****P= 5.88x10^−47^ (Welch’s t-test, Holm p-adjustment). In A and C, shading represents ± SEM.

### DEAF1 motifs are enriched at newly accessible regions near synaptic, but not early neuronal developmental genes

We hypothesized that the regulated binding of repressive factors at DARs might coordinate the repression of synaptic gene expression outside of developmental windows of synaptogenesis. To identify such factors, we searched for enriched TF binding motifs within DARs in a 3-kilobase window around each synaptic gene. This analysis revealed 21 enriched motifs, with the DEAF1 binding motif most significantly enriched (Figure 2B). DEAF1 is a highly conserved, neuronally enriched TF(*35–37*) that is associated with developmental delay and intellectual disability(*19–21*). DEAF1 directly binds DNA through its SAND domain and interacts with both positive and negative co-regulators through its MYND domain(*38*, *39*). Motif analysis within DARs around early neuronal development genes did not identify the DEAF1 binding motif (Figure S2A), indicating specificity at synaptic genes. Consistent with a specialized role, we observe a temporally regulated increase in accessibility across DEAF1 motifs at ∼10h embryogenesis that is specific to the nervous system(*25*) (Figure 2C).

### DEAF1 and CLAMP are predicted to co-repress synaptic gene expression

To independently identify candidate repressors, we took advantage of a third instar larval brain atlas of single-cell (sc) RNA-seq and scATAC-seq data(*40*) and the recently developed NetREm machine learning tool(*41*) (See Materials and Methods). As input we included 149 neuronal TFs identified from motif analyses (Figure S2A,Table S3). We first clustered wild-type cells from the scRNA-seq and scATAC-seq third instar larval brain atlas(*40*) broadly to capture substantial differences in synaptic gene expression during neuronal development into the following categories: neural progenitors, where synaptic genes are inactive; immature neurons, which express only low levels of synaptic genes; mature neurons, which broadly express synaptic genes; and glia, which were excluded from further analysis (Figure S2B-C). This analysis identified a candidate network of 10 TFs predicted to regulate at least 280 synaptic target genes, including DEAF1, which was predicted to negatively regulate 294 targets (60.6%; Figure S2D).

Together with the finding that the DEAF1 binding motif is highly enriched at newly accessible sites near synaptic genes, this prediction makes DEAF1 a strong candidate global repressor of synaptic gene expression. Therefore, we refocused the NetREm results to specifically identify a DEAF1 co-regulatory network (Figure 2D). Two TFs in the network have consistently high coordination scores (above |0.90|) and are predicted to act cooperatively with DEAF1 to negatively regulate synaptic gene expression: computed gene CG4360, a zinc-finger transcriptional repressor with multiple human orthologs(*42*), and Chromatin-linked adaptor for MSL proteins (CLAMP). CLAMP is a functional analog of mammalian CTCF that binds condensed chromatin and regulates the ability of other factors to bind DNA(*43*). Notably, our earlier motif analysis revealed that CLAMP motifs are the second-most enriched at newly accessible sites near synaptic genes (see Figure 2B) and CLAMP has been experimentally shown to physically interact with DEAF1(*44*). Together, these findings lead us to hypothesize that DEAF1 and CLAMP function together to repress synaptic gene expression outside of developmental windows of synaptogenesis.

Consistently, our analysis of third-instar larval ATAC-seq data indicates that DEAF1 and CLAMP binding motifs are co-accessible at a large fraction of synaptic genes. The DEAF1 binding motif is accessible at 328 synaptic genes (67.6%), the CLAMP binding motif is accessible at 300 synaptic genes (61.9%), and 239 synaptic genes (49.3%) have accessible motifs for both DEAF1 and CLAMP (Figure 2E). Of these 239 synaptic genes, 105 have spatially neighboring co-accessible motifs (Table S4, see Materials and Methods). Measured separately, the accessibility of DEAF1 and CLAMP motifs are respectively 4-fold and 3.5-fold more enriched at synaptic than at random nervous system genes (Figure 2F, padj = 3.5x10^-48^ and 1.1x10^-39^). Notably, accessibility of both motifs is even more significantly enriched at synaptic genes (Figure 2F, padj = 1.0x10^-52^), supporting the hypothesis that DEAF1 and CLAMP function together to regulate synaptic gene expression.

The DNA binding domain of DEAF1 is highly conserved in mammals and previous studies have shown that DEAF1 bidirectionally regulates transcription of the serotonin receptor 5-HT1A, encoded by the *HTR1A* gene(*45*, *46*). Notably, there is a DEAF1 binding site at both human and *Drosophila* HTR1A (Figure 2G-H) We hypothesized that DEAF1 may similarly regulate additional coexpressed synaptic genes in humans. In data from human forebrain organoid derived neurons, we identified the DEAF1 binding motif in annotated peaks of accessible chromatin at highly ranked orthologs of *Drosophila* synaptic genes, including *brp/ERC1*, *mAChR-A/CHRM3*, and Syt1/*SYT1* (Figure 2I-J, S2E-H). To investigate broader DEAF1 motif enrichment at synaptic genes, we identified genes with expression patterns closely correlated (> 0.8 Spearman) to the average expression profile of synaptic footprint orthologs in time-series data from the human developing brain(*47*). We then examined accessible chromatin regions at these coexpressed genes in ATAC-seq data from human forebrain organoid derived neurons(*48*) and found significant enrichment of the DEAF1 motif compared to random nervous system genes (Figure 2K). These data suggest that broad co-regulation of synaptic genes by DEAF1 may be conserved in mammals.

### *deaf1* or *clamp* knockdown de-represses synaptic gene expression

Based on these analyses, we hypothesize that DEAF1 and CLAMP function together in the coordination of synaptic gene expression across the brain by broadly repressing synaptic gene expression outside of windows of peak synaptogenesis. If this model is correct, loss of either factor would be expected to dysregulate synaptic gene expression. To test our model *in vivo*, we focused on the third instar larval stage when synaptic gene expression is normally repressed. We used *elav*-Gal4, which drives expression in all neurons and salivary glands, to express RNAi constructs targeting *deaf1* or *clamp* and confirmed knockdown of each factor on chromatin (Figure S3A-D). We then conducted RNA-seq on isolated larval brains.

Synaptic genes are significantly enriched in the upregulated groups of genes in both *deaf1* (odds.ratio = 6.28, padj = 1.68x10^-68^) and *clamp* (odds.ratio = 4.44, padj = 1.09x10^-28^) knockdowns. The overlap of these differentially upregulated genes confirm significant intersection among synaptic genes upregulated with either TF knockdown (odds.ratio = 6.98, padj = 5.22x10^-27^). In contrast, there is no enrichment for synaptic genes in downregulated genes (Figure 3A).

**Figure 3.**
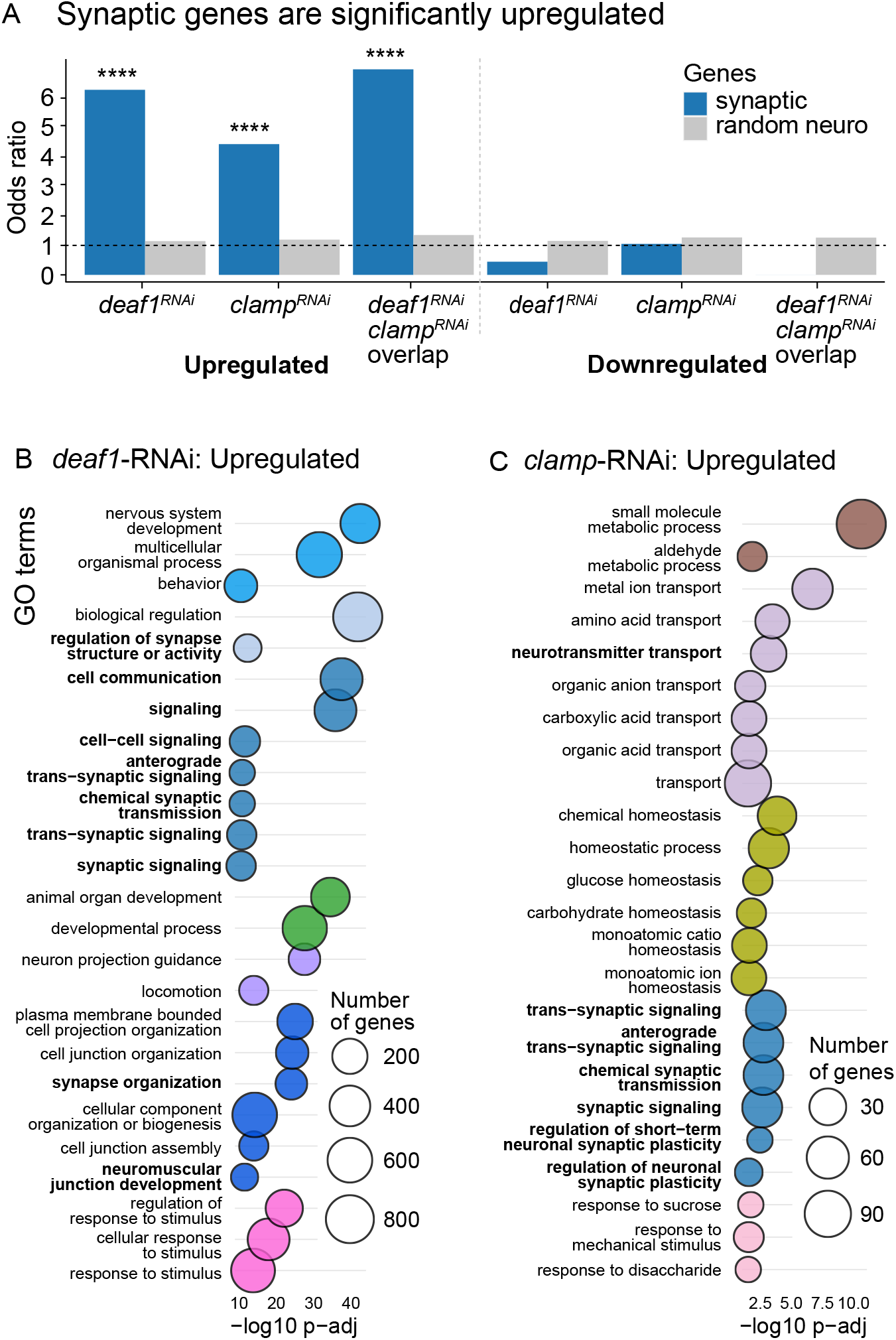
*deaf1* or *clamp* knockdown de-represses synaptic gene expression. (A) Odds ratios of genes differentially regulated in third instar larval brains upon knockdown of *deaf1* or *clamp*. Synaptic genes (blue) are significantly enriched among upregulated genes, but not among downregulated genes. Random neural genes (grey) are not enriched among up or downregulated genes. (B-C) GO analysis of genes upregulated upon knockdown of *deaf1* (B) or *clamp* (C) show enrichment for synaptic processes (bold). GO terms are grouped by hierarchical parent terms and trimmed for biological process terms with uniqueness scores > 0.9, showing the top 25 terms for (B) and all terms for (C). GO terms are color coded based on shared hierarchical parent GO terms. In A, ****P<10^-5^ (Fisher’s exact test, Bonferroni p-adjustment).

Knockdown of *deaf1* resulted in the differential expression of 2862 genes relative to RNAi controls: 1558 upregulated and 1304 downregulated. GO analysis of genes significantly upregulated following *deaf1* knockdown revealed enrichment for cell communication terms, including regulation of synapse structure or activity and synaptic signaling (Figure 3B, bold). Consistent with broad DEAF1-mediated repression of synaptic genes, all of the top 15 GO terms from the 485 coordinately regulated synaptic genes (Figure 1C) are enriched across the significantly upregulated genes. In contrast, GO enrichment analysis of genes downregulated upon *deaf1* knockdown yielded broad metabolic process and DNA replication terms (Figure S4A).

Out of 485 coordinately regulated synaptic genes, 207 are upregulated upon *deaf1* knockdown, while only 23 are downregulated. Among the upregulated genes are those that encode vesicular and active zone components, including nSyb, Brp, Rim, Syt1, Ca-β, synaptic cell adhesion molecules, including multiple Dpr, Side, Beat, Nlg, CadN proteins, and neurotransmitter transporters and receptors, including vesicular glutamate and monoamine transporters VGlut and Vmat, choline acetyltransferase (ChAT) and receptors for acetylcholine, GABA, glutamate, dopamine. The serotonin receptors 5-HT1A and 5-HT1B were also upregulated, indicating conserved roles for DEAF1 in *Drosophila* and mammals(*45*, *46*). Overall, these data confirm that DEAF1 is a broad repressor of pan-neuronal and subtype-specific synaptic gene expression.

Knockdown of *clamp* in all neurons resulted in the differential expression of 1566 genes relative to RNAi controls, 861 upregulated and 703 downregulated. Similar to *deaf1* knockdown, of 485 co-regulated synaptic genes, 106 were upregulated and only 27 downregulated. GO analysis of all upregulated genes shows enrichment for neurotransmitter transport and synaptic signaling terms (Figure 3C, bold). We also observe GO term enrichment of upregulated genes for small molecule metabolism and amino acid transport (Figure 3C, brown, purple), suggesting that CLAMP has additional broader roles in negatively regulating gene expression in the nervous system consistent with our prior findings(*49*). Downregulated genes are enriched for protein catabolic process and endocytosis terms (Figure S4B). Synaptic genes upregulated upon *clamp* knockdown include 63 genes also upregulated when *deaf1* is knocked down, including genes encoding both pan-neuronal synaptic proteins such as Syt1 and subtype-type specific synaptic proteins such as Vmat, ChAT, and receptors for acetylcholine, glutamate, and dopamine. We observe a similar upregulation of synaptic genes in *clamp* null third instar larval brains(*49*). Together, these findings demonstrate that disruption of *deaf1* or *clamp* broadly de-represses synaptic gene expression in the late larval nervous system.

### DEAF1 and CLAMP limit synapse formation

Synapse formation and refinement are critical developmental processes. In wild-type animals synaptic gene transcription peaks during late nervous system development, then falls to low levels following this window of robust synaptogenesis (see Figure 1A). This observation suggests that broadly coordinated activation and repression of synaptic genes act together to regulate proper wiring of the nervous system.

To determine the role of DEAF1/CLAMP-dependent synaptic gene repression in synaptogenesis, we turned to the well-characterized, highly stereotyped third instar larval neuromuscular junction. Here, glutamatergic motor neurons innervate single muscle cells and form a stereotyped number of synaptic boutons, each containing ∼10 individual synapses, allowing us to readily quantify synapse formation in different genetic backgrounds. Pan-neuronal knockdown of *deaf1* caused a 56% increase in synaptic bouton number and significantly increased single synapse number compared to control (Figure 4A-E). We also observe a 54% increase in synaptic boutons in a heteroallelic combination of *deaf1* alleles (*deaf1^k3/S10B^*) that produces rare escapers that survive to third instar larvae (Figure S5A-C). Similarly, pan-neuronal knockdown of *clamp* resulted in a 28% increase in synaptic bouton formation and significantly increased synapse number (Figure 4A-E). In *clamp^KG00473^* hypomorphs, we observe a 24% increase in synaptic boutons (Figure S5D-F), similar to previous observations(*50*). Thus, transcriptional repression of synaptic gene expression by DEAF1 and CLAMP is required to constrain synapse formation in the developing nervous system.

**Figure 4.**
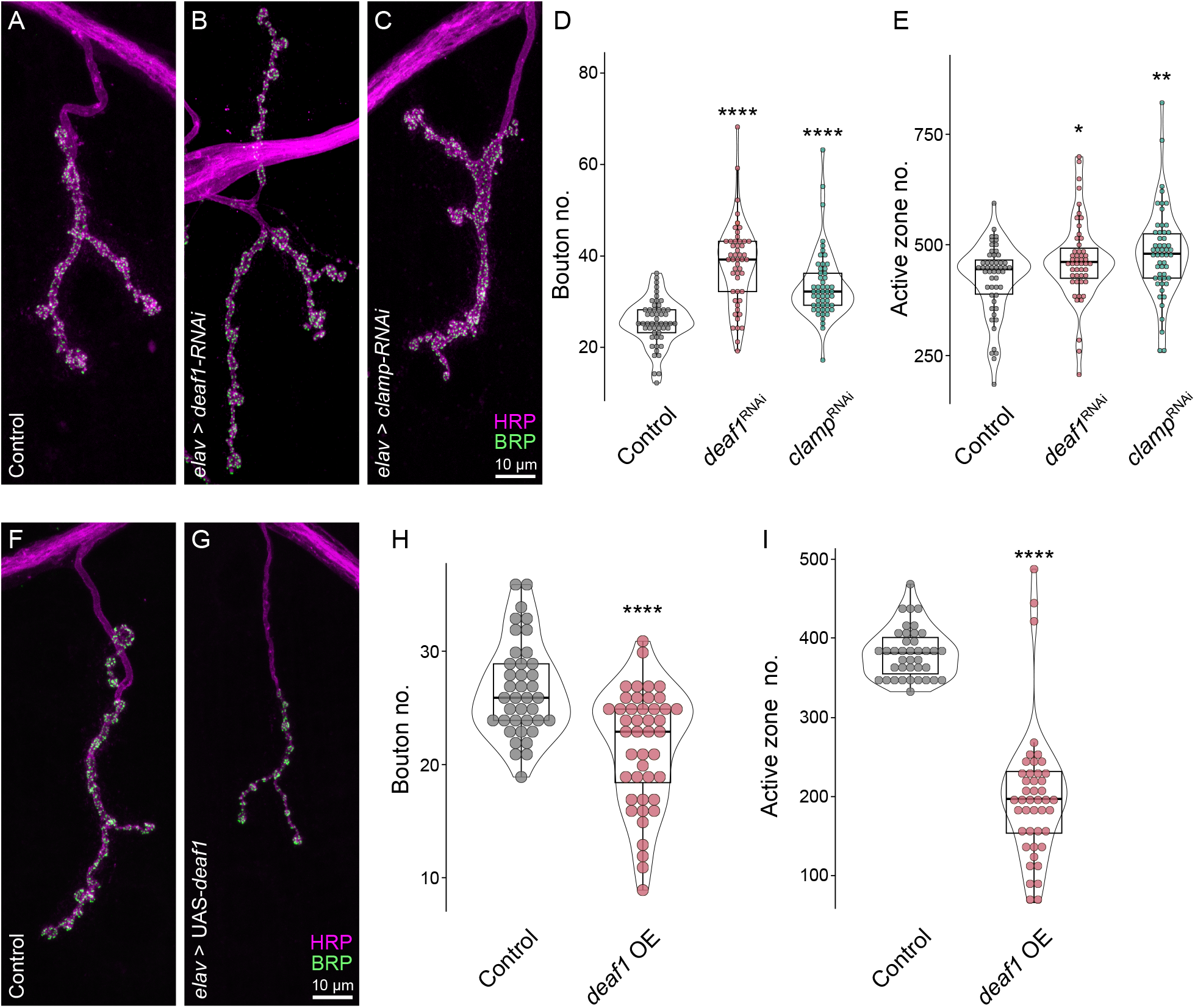
DEAF1 and CLAMP limit synapse formation. (A-E) Representative images (A-C) of NMJ 4 in control (*gfpRNAi*) (A), *deaf1RNAi* (B), *clampRNAi* (C), and quantification of synaptic bouton number (D) and active zone number (E). (F-I) Representative images of NMJ 4 in control (F), *deaf1* overexpression (G) and quantification of synaptic bouton number (H) and active zone number (I). Scale bar = 10 μm. Data points represent individual NMJs. Kruskal-Wallis test followed by Dunn’s multiple comparisons test and Benjamini-Hochberg p-value adjustment. In D, E, H, and I, ****padj<10^-6^, **<0.01, *<0.05. Data is represented as mean ± SEM.

These findings suggest that activation and repression of synaptic gene transcription is dynamically controlled through the temporally regulated actions of TFs acting in opposing directions to regulate synapse formation. This further suggests that dysregulated repression of synaptic genes might inhibit the formation of synaptic connections. To determine whether DEAF1 misexpression is sufficient to repress synapse formation, we overexpressed *deaf1* in developing neurons and quantified synapses at the larval NMJ. We observed a 19% decrease in bouton formation (Figure 4F-H) and a striking 52% decrease in single synapses (Figure 4I). Thus, the dynamic regulation of synaptic gene activation and repression plays a critical role in sculpting neuronal connectivity during nervous system development.

### DEAF1 directly binds and represses synaptic genes in a CLAMP-dependent manner

We next sought to determine if DEAF1 and/or CLAMP directly occupy synaptic gene regulatory regions to repress their transcription. As a pioneer TF, CLAMP can increase chromatin accessibility(*22*, *43*) and direct distinct TFs and chromatin regulators to the correct genomic locations to drive context-specific functions, including directing dosage compensation(*51*), splicing(*52*), and polymerase pausing(*23*). Therefore, we hypothesized that CLAMP might promote the recruitment of DEAF1 to facilitate repression of synaptic genes outside of developmental windows of synaptogenesis. Consistently, when we co-label DEAF1 and CLAMP on polytene chromosomes, we observe colocalization at a subset of sites across all chromosomes (Figure S6A).

To directly test if DEAF1 occupies synaptic gene regulatory regions during the repressed state, we profiled DEAF1 binding by CUT&RUN(*53*, *54*) in third instar larval brains, when synaptic genes are transcriptionally repressed. In RNAi control brains, we found that DEAF1 occupancy is strongly enriched at both DARs and transcription start sites (TSSs) of synaptic genes relative to random nervous system genes (Figure 5A–B, red vs. gray). At DARs associated with early neuronal developmental genes, DEAF1 occupancy was detected at levels closer to random nervous system genes (Figure S6B). In contrast, IgG controls showed minimal enrichment across all three gene sets (Figure S6C), supporting specific association of DEAF1 with synaptic regulatory regions. Peak-centered heatmaps further showed strong DEAF1 enrichment at called peak summits relative to the IgG negative control (Figure S6D). Consistent with this pattern, DEAF1 binding across temporal expression clusters was particularly prominent in clusters 8, 2, and 1, with the strongest enrichment observed in cluster 8, which is enriched for coordinately regulated synaptic genes (Figure S7A; see Figure 1D, E). Thus, DEAF1 directly occupies synaptic gene regulatory regions during the larval repressed state.

**Figure 5.**
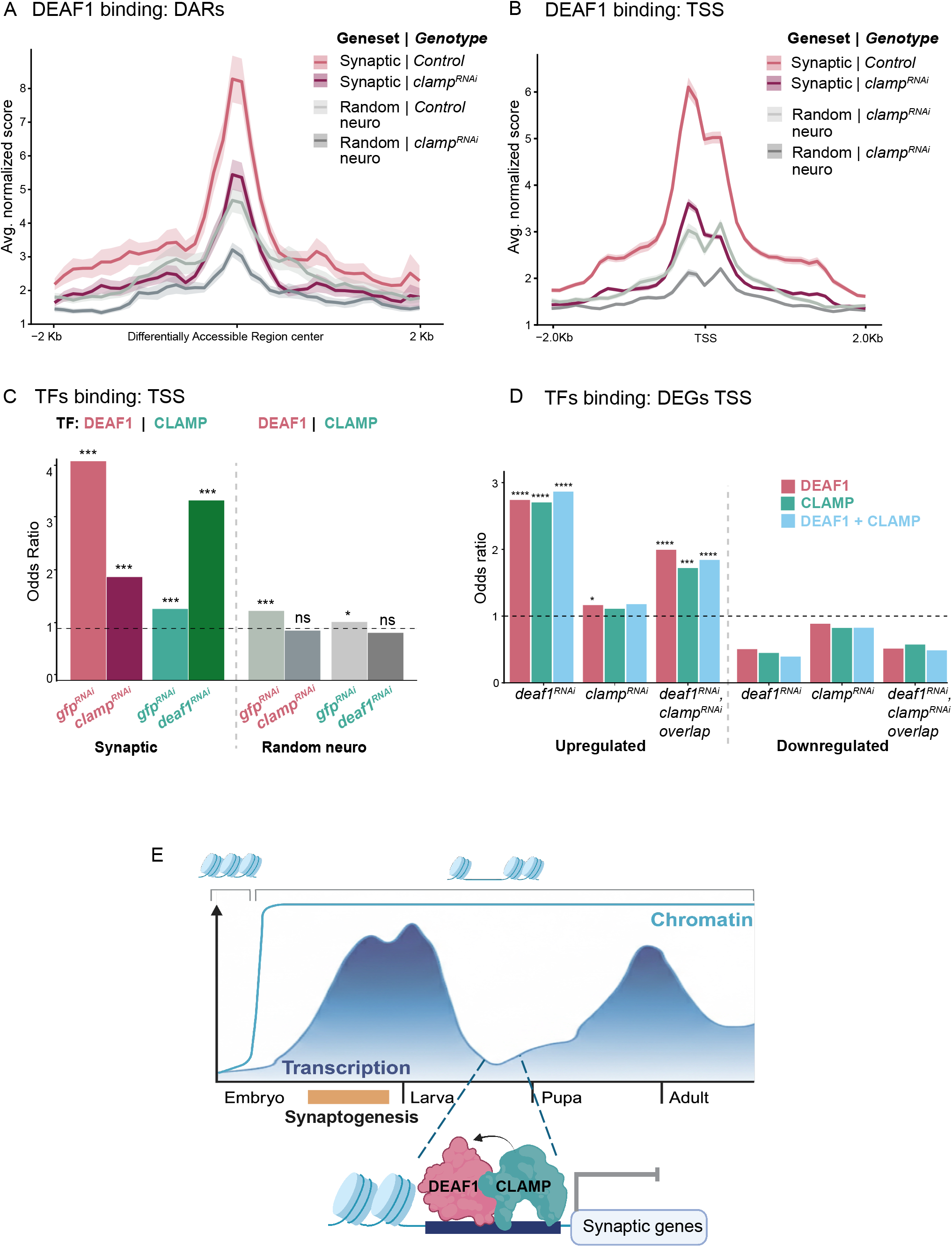
DEAF1 directly binds synaptic DARs in a CLAMP-dependent manner and represses synaptic genes. (A) Average DEAF1 CUT&RUN enrichment across synaptic DARs (control, dark maroon) and random nervous system genes (grey) in control (*gfpRNAi*) and *clampRNAi* third instar larval brains. (B) Average DEAF1 CUT&RUN enrichment across synaptic promoter regions (control, dark maroon) and random nervous system genes (grey) in control (*gfpRNAi*) and *clampRNAi* third instar larval brains. (C) Odds ratios of DEAF1 binding at synaptic vs. random nervous system genes in control (*gfpRNAi*) and *clampRNAi*. (D) Odds ratios of DEAF1 binding at synaptic and random nervous system genes differentially up-or downregulated upon *deaf1RNAi*, *clampRNAi*, or in both (overlap). In C, D ****padj<10⁻⁵, ***<0.001, *<0.05, ns (Fisher’s exact test, Bonferroni adjustment). Shading represents ± SEM. (E) Model for temporal regulation of synaptic gene transcription by CLAMP and DEAF1. Chromatin accessibility increases across synaptic gene regulatory regions preceding synaptic gene expression and peak synaptogenesis during embryonic nervous system development. Following synaptogenesis, these regions remain accessible while synaptic gene transcription is repressed. CLAMP promotes DEAF1 occupancy at accessible synaptic regulatory regions during this repressive state, thereby restricting synaptic gene expression and limiting premature or excessive synapse formation.

To test the model that CLAMP recruits DEAF1, we profiled DEAf1 occupancy following knockdown of *clamp* in the nervous system, again using *elav*-GAL4 driven RNAi. When *clamp* is knocked down, DEAF1 occupancy at synaptic DARs is substantially reduced relative to control (Figure 5A-B, red vs. magenta), indicating that CLAMP is required to promote and/or stabilize DEAF1 binding at these sites. As expected for a pioneer transcription factor that opens chromatin at many locations genome wide(*43*), CLAMP also promotes chromatin opening at random nervous system genes (Figure 5A-B, dark vs. light gray). Consistently, CLAMP CUT&RUN confirmed CLAMP occupancy at neuronal DARs and promoters, including those associated with synaptic genes (Figure S7B-C).

To determine if DEAF1 chromatin binding is associated with direct transcriptional regulation, we next compared TSS-associated DEAF1 CUT&RUN occupancy with genes differentially expressed following *deaf1* and *clamp* knockdown. DEAF1 binding was significantly enriched at synaptic genes upregulated upon *deaf1* knockdown and at synaptic genes upregulated in both *deaf1* and *clamp* knockdowns, but not at downregulated genes (Figure 5C), consistent with DEAF1 functioning primarily as a repressor at these loci. Finally, to define high confidence direct targets, we investigated the intersection between DEAF1-bound synaptic genes, genes containing a differentially accessible DEAF1 motif, and genes upregulated upon *deaf1* knockdown. Of the 207 synaptic genes upregulated upon *deaf1* knockdown, 190 (92%) were directly bound by DEAF1 (Figure 5D). 147 of these synaptic genes also contained a differentially accessible DEAF1 motif, strongly supporting the conclusion that the majority of de-repressed synaptic genes are direct DEAF1 targets.

Collectively these findings indicate that DEAF1 directly occupies DARs at synaptic genes to repress transcription following embryonic nervous system development, and that CLAMP is required to promote DEAF1 binding at these loci. Combined with the persistence of chromatin accessibility across later developmental stages, these data support a model in which temporal repression of synaptic genes is achieved not by chromatin closure, but through CLAMP-dependent recruitment of the DEAF1 repressor to regulatory elements that remain accessible and poised for subsequent activation(Figure 5E). In this model, accessible chromatin is established early and maintained across development, while DEAF1–CLAMP occupancy provides a temporally restricted repressive state that prevents premature synaptic gene expression until the appropriate developmental window. Together, our gene expression, synaptic imaging, and transcription factor occupancy data underscore the critical role of repression in coordinating gene expression during circuit development.

## Discussion

Combining multimodal genomic analysis with predictive modeling and *in vivo* loss-and gain-of-function validation, we reveal a central role for the coordinated repression of synaptic genes in controlling synaptogenesis and identify DEAF1 and CLAMP as key repressors.

Neural circuitry is tightly controlled through the regulation of synapse formation, plasticity, and maintenance. Altered synapse formation and circuit function are common features of neurodevelopmental disorders(*55*, *56*), highlighting the importance of proper regulation. The formation of functional synapses necessitates the coordination of hundreds of proteins across pre-and postsynaptic cells, suggesting that synaptic gene expression must be coordinated across neuronal subtypes. Indeed, we find that large numbers of synaptic genes, including those encoding pan-neuronal and subtype-specific synaptic components, are coordinately activated and repressed at the level of transcription in the *Drosophila* nervous system. This is consistent with observations from a time-series single-cell transcriptomic study in the *Drosophila* visual system, which identified ∼200 genes that are temporally coordinated across at least 50% of cell types and highly enriched for synaptic genes(*29*). Notably, a recent study found that neuronal transcriptomes become temporally synchronized over the course of maturation, with the transcriptomes of early and late-born neurons becoming indistinguishable following the axon pathfinding/target recognition period when transcriptional diversity peaks(*57*). Our analyses suggest that the coordinated opening of chromatin across pan-neuronal and subtype-specific synaptic genes immediately preceding synapse formation acts as a permissive brain-wide signal that synchronizes the activation of synaptic gene expression across neuronal subtypes.

While synaptic gene expression is also coordinatedly repressed outside developmental windows of peak synaptogenesis, these synaptic DARs remain open during these periods, indicating that repression occurs through a distinct mechanism. We identified DEAF1 as the TF most associated with synaptic DARs and a predicted repressor. DEAF1 is a multifunctional TF that interacts with an array of co-regulators that both positively and negatively regulate transcription(*58–60*). Through GRN analysis, we identified CLAMP, which has also been shown to positively and negatively regulate transcription, as a candidate DEAF1 coregulator. Notably, CLAMP motifs are the second-most enriched at dynamically regulated sites upstream of synaptic genes. We confirm DEAF1 and CLAMP as negative regulators of synaptic gene expression and demonstrate that the broad repression of synaptic gene expression plays an instructive role in constraining synaptic connectivity: Loss of either DEAF1 or CLAMP results in excess synapse formation. Conversely, neuronal overexpression of DEAF1 blocks synapse formation.

Consistently, we observe DEAF1 and CLAMP colocalizing on polytene chromosomes and DEAF1 and CLAMP motifs are co-accessible at a large fraction of synaptic genes. Furthermore, we find that DEAF1 is directly bound at genes upregulated in the absence of each factor but not at downregulated genes, demonstrating a direct role in repressing synaptic genes. Both pan-synaptic and subtype-specific synaptic genes are upregulated and bound by DEAF1, suggesting DEAF1 temporally coordinates synaptic gene expression across neuronal subtypes during nervous system development, consistent with its broad neuronal expression.

Notably, DEAF1 occupancy at synaptic DARs is substantially reduced upon *clamp* knockdown. Together with the reported physical interaction between DEAF1 and CLAMP(*44*) and their co-accessible motifs, this dependence supports a model in which CLAMP facilitates recruitment of the DEAF1 repressor to synaptic DARs, rather than the two factors binding independently. How CLAMP promotes DEAF1 binding remains an open question. As a GA-rich pioneer factor, CLAMP increases chromatin accessibility and recruits diverse partners, including the MSL dosage compensation complex, to specific genomic locations(*43*, *51*, *61*, *62*). Because the DARs bound by DEAF1 remain accessible throughout the stages we examined, our data suggest that at synaptic genes CLAMP may act less to open chromatin than to guide or stabilize DEAF1 occupancy on regulatory elements that are already accessible. Determining whether CLAMP recruits DEAF1 through direct interaction, through a shared partner, or by locally shaping chromatin at these sites will be an important direction for future work.

The neurotransmitters released and interpreted by neurons are a defining part of their identity. It has been well established that neuronal identity is conferred by cell-specific combinations of TFs termed terminal selectors(*4*, *14–16*). In contrast, pan-neuronal gene expression appears to be activated in a cell-specific manner by a diversity of redundant factors(*17*, *18*, *63*). Together with our findings, this suggests a regulatory logic in which activation of coordinated synaptic gene expression relies on the combined action of dynamic chromatin accessibility, terminal selectors, and distinct combinations of redundant TFs while repression is controlled by repressive TFs that act broadly across cell types. Combined, these activating and repressive, local and global regulatory mechanisms can coordinate the acquisition of distinct neuronal fates with circuit assembly.

We hypothesize that maintaining chromatin accessibility at synaptic genes following developmental windows of synaptogenesis facilitates synaptic plasticity in the brain by allowing for the rapid activity-dependent transcriptional activation of synaptic genes. A key question to be addressed in the future is how the transitions between activation and repression of synaptic genes are regulated. One possibility is the dynamic regulation of DEAF1 and/or CLAMP themselves. DEAF1 can both positively and negatively regulate transcription of the serotonin receptor 5-HT1A with GSK3-dependent phosphorylation promoting its repressive role(*45*, *46*). In fact, recent work suggests that DEAF1 promotes activation of axon guidance genes in an organoid model (*64*). The same transcription factors often regulate either activation or repression at different times and genomic locations depending on partner proteins and post translational modifications(*65*). Therefore, it is possible that DEAF1 remains at its binding sites during both activation and repression, with its function modulated by phosphorylation state. It is also possible that GAF(*66*), a well studied transcriptional activator, replaces the repressive CLAMP during gene activation because they can compete for very similar binding sites(*61*). Both models will be important to test in the future.

One mechanism by which the modulation of DEAF1 through phosphorylation and/or the replacement of CLAMP with GAF might control synaptic gene expression is the regulation of RNA Pol II 5’ end pause release. We found that differentially accessible DEAF1 binding sites at synaptic genes are enriched at 5′ ends, consistent with a potential role in regulating Pol II pausing and the binding of mammalian DEAF1 at 5′ ends of several candidate synaptic genes(*36*, *45*, *46*). Future analysis of RNA Pol II pausing will be required to determine whether synaptic gene expression is regulated by pause release.

Our findings reveal the importance of regulated repression in sculpting synaptic connections and provide a mechanistic framework for diverse regulatory mechanisms acting together to dynamically modulate synaptic gene expression. Both dominant and recessive pathogenic *DEAF1* alleles cause syndromic intellectual disability(*19*, *20*), underscoring the importance of decoding the regulatory logic of coordinated synaptic gene expression in neurodevelopment.

## Materials and Methods

### Drosophila stocks

The following fly lines used in this study were obtained from the Bloomington Drosophila Stock Center (BDSC, NIH P40OD018537): *w^1118^* (RRID:BDSC_5905), *UAS-deaf1-RNAi* (RRID:BDSC_32512), *deaf1^k3^* (RRID:BDSC_26279), *deaf1^S10B^*(RRID:BDSC_91626), *Clamp^KG00473^* (RRID:BDSC_14352), *UAS-GFP-RNAi* (RRID:BDSC_9330), *pros-Gal4* (RRID:BDSC_80572), *elav^C155^-Gal4* (RRID:BDSC_458). Additional fly lines used in this study were obtained from The Zurich ORFeome Project (FlyORF)(*67*): UAS-*deaf1-3xHA* (RRID:Flybase_FBst0501296). *Drosophila melanogaster* stocks were raised on molasses food (Lab Express, Type R) in a 25°C incubator with controlled humidity and 12h light/dark cycle.

### Temporal gene expression profile analysis

Transcriptomic data was accessed through FlyBase (release FB2020_05)(*9*, *42*) from Graveley et al., 2011, *Nature*. Data was separated by developmental timepoints. Embryonic 0-2h through pupal p15 developmental timepoints were used for finding Spearman correlations to gene sets with the average profile of 20 synaptic genes using cor() in the R stats package(*68*). For the early neuronal developmental geneset, 46 genes with Spearman correlation scores above 0.71 were not found in nervous system expression data and were not included in the final list of 485 genes. Random nervous system expressed control gene sets were generated from genes expressed in the nervous system with Spearman correlations to synaptic genes below 0.80 and with sample_n() in the dplyr package(*69*).

#### Clustering analysis

Kmeans clustering was performed with the R stats package. The number of clusters was determined from the gap statistic calculated with packages cluster(*70*) and factoextra(*71*). Gene ontology analysis was performed with the R package gprofiler2(*72*) and sorted into parent terms with rrvgo(*73*).

#### Single-cell temporal profile analysis

Using Seurat(*74*), all nervous system annotated clusters were subset from 2h embryonic datasets(*25*), as Seurat objects, then merged. Larval nervous system datasets(*26*) and pupal visual system datasets(*29*) from each timepoint were loaded as count matrices. Seurat objects were created and filtered with min.cells = 3, min.features = 200, percent.mt < 5, then merged. Each merged object was normalized, 5000 variable genes were used for principle component analysis (PCA) and 50 PCs were used to identify nearest neighbors and RunUMAP with clustering resolutions of 0.47 for embryo and 0.6 for larva and pupa. The R package scCustomize(*75*) was used with AddModuleScore to average the expression of all synaptic genes in our list for each cell, and the score of each cell was averaged per timepoint.

### Chromatin accessibility analysis

DNase-seq bigwig files were downloaded from the Furlong laboratory website(*31*), and ATAC-seq bigwig files from NCBI GEO GSE263159(*33*) and GSE226514(*34*). Average profiles of accessibility at synaptic genes were made using deepTools(*76*) computeMatrix with parameters ‘computeMatrix reference-point --referencePoint TSS -a 1000 -b 1000 --binSize 100 --skipZeros --smartLabels --sortRegions descend’ and plotProfile ‘--plotType se --yMin 0 --yMax 350’’. To bring scores per genomic region into similar ranges across different laboratories and experiment types, a scale factor was applied for each: DNase-seq embryos (1/100), GSE263159 (1), and GSE226514 (1/25).

### Protein expression data analysis

Protein expression data from Casas-Vila et al., 2017(*32*) (Table S1). The imputed.log2.LFQ.intensity data for each protein were averaged across the four samples at each timepoint. The data were then subset to synaptic proteins (n=19), scores were shifted by the minimum average imputed score, and the score of each gene normalized to its maximum expression value.

### Motif discovery

Bigwig files were converted to bedgraph format with bigWigToBedGraph(*77*). Peaks were called with MACS2(*78*) bdgpeakcall with a cutoff above the standard deviation and parameters ‘--max-gap 10 --min-length 150’. The neuronal 10-12h peak file was subset using BEDTools(*79*) ‘subtract -A’ to remove accessibility peaks intersecting those present in neuronal 8-10h. Likewise neuronal 10-12h peaks were removed from neuronal 8-10h peaks to use as background control regions in motif analysis. Peaks newly accessible in the neuronal 10-12h data were subset to those within 3kb of synaptic genes or within 3kb of early neuronal development genes using bedtools ‘window -w 3000 -u’. Homer(*80*) findMotifGenome.pl was used for motif analysis with the parameters ‘dm3 -len 8,10,12 -size given -p 16 -dumpFasta’. The newly accessible neuronal 10-12h peak regions were used as input with the peaks unique to neuronal 8-10h data as background controls.

### Single-cell GRN analysis

#### Single-cell data processing for NetREm input

Third instar larval brain scRNA-seq and scATAC-seq data(*40*) were processed with Seurat(*74*) and Signac(*81*), objects were created from counts matrix then filtered, normalized, and clustered using the parameters detailed in Mohana et al.(*40*). Genes with highest average expression in each scRNA cluster (gene activity scores for scATAC), were used with enriched marker genes from FindAllMarkers(seurat_obj, only.pos = TRUE, min.pct = 0.25, logfc.threshold = 0.25, test.use = “wilcox”) to broadly annotate the 16 RNA and 19 ATAC clusters, which were then merged by annotation.

#### NetREm

NetREm(*41*) is a tool that predicts TF-target gene (TG) regulatory networks (complementary GRN) and TF-TF coordination networks (at the TG-level and overall) for the cell-type in a given context. The TF-TG regulatory network scores TFs by their ability to predict TG expression levels. The TG-specific TF-TF coordination B network analyzes how TFs coordinate antagonistically or cooperatively to co-regulate the given TG. The overall TF-TF coordination network, B^_, summarizes the net behavior among TFs to co-regulate TGs overall. As input, NetREm requires gene expression data (single-cell or bulk-level) and a prior protein-protein interaction (PPI) network among the predicted TFs. We applied NetREm according to the methods described in Khullar et al., 2025(*41*), with the following specifications.

For NetREm TF input, we included 149 TFs identified as best matches and lower affinity score matches from motif discovery analyses. Of these, 135 were identified in the third instar larval brain scRNA-seq expression data. For TGs we included all 485 genes from our synaptic genes list of which 457 were identified in the larval data. For PPI Network data, we used the full network of combined scores from *Drosophila melanogaster* STRINGdb version 12(*82*). STRINGdb v12 was subset to include 2,046 unique TF-TF edges of known direct and/or indirect PPIs, comprising 133 of the 135 default TFs examined in this analysis, E(spl)mbeta-HLH and E(spl)mgamma-HLH were not identified in the PPI data and given artificial edge weights of 0.01.

We derive prior reference GRN information that predicts TFs that associate to regulatory elements to regulate TGs. To do this, we utilize various available multiomics data to identify candidate TFs that may bind to regions of open chromatin linked to those TGs. For TG accessibility regions, we identified all chromatin accessibility peaks in the third instar larval brain scATAC-seq data that were found within 3kb of synaptic genes and subset them according to broad cell-type annotations. We performed motif analysis on these accessible regions using motifmatchr(*83*) matchMotifs function with the JASPAR2024 motif database and BSgenome.Dmelanogaster.UCSC.dm6 genome and output scores for each peak. We retained the top 50% TFs for each peak based on the highest scores. Only 96 of the 135 TFs were identified by JASPAR2024; thus, we queried additional motif databases using query(MotifDb(*84*), andStrings=c(”Dmelanogaster”), orStrings=c(”jaspar2024”, “Dmelanogaster”), notStrings=”ctcfl”). These additional motifs are drawn from Cisbp_1.02, FlyFactorSurvey, JASPAR_2014, JASPAR_CORE, Jaspar2016, Jaspar2018, and Jaspar2022. To identify the motifs in the accessibility sequences, we used monaLisa findMotifHits function with parameters ‘query=pwm_list, subject=seqs, min.score=”75”, method=”matchPWM”, BPPARAM=BiocParallel::SerialParam()’. We found 112 of the 135 TFs using MonaLisa (min. score percentile 75), with 91 TFs in common with the JASPAR2024 database for a total of 117 TFs with at least one instance of accessible motif. We thus derive our motif-based reference GRN for these TGs in cell-types that have identified accessible regions or motifs.

For the TGs in our motif-based reference GRN, we input the TG-specific cell-type TFs to NetREm as their respective candidate TFs. For the remaining TGs (with unmapped prior GRN information), we use the 135 default TFs; these include 131 in glia, 74 in neural progenitors, 140 in immature neurons, and 97 TGs in mature neurons. Since few synaptic genes are expressed in glial cells, glial analysis was removed from final comparisons. Overall, we run NetREm step-by-step for each of these TGs for their corresponding set of TG-specific TFs (if TG is in motif-based reference GRN) or default TFs (if TG is not in reference GRN).

We ran NetREm step-by-step for each TG with the following parameters. We randomly selected 80% of the samples in the gene expression data for training (M cells) the model and the remaining 20% for testing. We selected β = 100 for our PPI network prior constraint to stress identifying groups of functionally coordinating TFs for regulating the TG, and emphasizing known direct and/or indirect PPIs among the TFs. Potential indirect PPIs are not found in *Drosophila melanogaster* STRING v12 database after subsetting for direct, physical PPIs We trained the model on the training gene expression data. The sparsity parameter α is selected using LassoCV with 5-fold cross-validation to optimize the mean square error on training data. We do not fit any y-intercept.

### DEAF1 and CLAMP motif analysis

From third instar larval brain ATAC-seq data(*33*), regions within 3kb of synaptic genes and random nervous system gene sets were identified using BEDtools ‘window -w 3000’. We generated a GRanges object of peak positions and used the MotifDb(*84*) R package to call Position Frequency Matrices (PFMs) for DEAF1 and CLAMP binding motifs from the JASPAR2022 database. PFMs were converted to objects of class PWMatrix from the TFBSTools package using the convert_motifs() function from the universalmotif package. Motif matching was performed using the matchMotifs() function from motifmatchr(*83*) with p.cutoff = 5e-05, genome = “dm6”, out’ = ‘positions’. This generates the coordinates of all hits for each motif in our database. Odds ratios and Fisher’s exact test significance scores were calculated with the R package GeneOverlap/1.36.0(*85*).

#### Human motif analysis

Human orthologs of the 20 *Drosophila* synaptic footprint genes that are annotated as high or moderate were retrieved from the DRSC integrative ortholog prediction tools(*86*). Processed temporal single-cell brain data from developing human brain(*47*) was downloaded from https://github.com/linnarsson-lab/developing-human-brain/ in .h5ad format. We used normalized and scaled counts to calculate the average expression for each gene, region, and timepoint. Regions with only one timepoint were removed. For each gene across region and timepoint, we calculated the Spearman correlation distance from the average expression pattern of the synaptic orthologs.

Gene locations were retrieved from the GRCh38.114 genome for all genes above 0.8 Spearman correlation to expression pattern and an equal number of random nervous system expressed genes from less than 0.8 Spearman correlation. Peak files of temporally accessible chromatin from human forebrain organoids were downloaded from NCBI GEO GSE132403(*48*). Neuronal ATAC-seq datasets were subset to include peaks within 3kb of synaptic correlation or random nervous system gene sets. The R packages MotifDb, universalmotif, and motifmatch were used to query the DEAF1 binding motif from Jaspar2022, convert to Position Weight Matrix format, and search the subset accessibility peaks for motif presence.

### Polytene chromosome immunostaining and analysis

Polytene chromosome squashes were prepared from third instar salivary glands of control (*elav>GFP RNAi*) and *deaf1* and *clamp* knockdown (*elav>deaf1 RNAi, elav>clamp RNAi*) genotypes using the protocol described in Rieder et al.(*87*). For immunostaining, we used rabbit anti-CLAMP (1:500, SDIX) and guinea-pig anti-DEAF1 (1:500, gift from William McGinnis(*88*)) antibodies and species specific Alexa Fluor secondary antibodies at 1:500. All samples were then washed in PTX, stained with DAPI, and mounted in ProLong Diamond (Thermo Fisher Scientific). We imaged slides at 40X on a Nikon Spinning Disc Confocal Microscope. Polytene images were analyzed using NIS Elements. Histone loci were identified by Clamp staining and a 70um line was drawn from the center of the histone locus down the chromosome using the Profile Analysis Tool. Line profile fluorescence intensity was measured in each channel for the region including a neighborhood of ∼1um on either side of the line. Intensities were compared by Kruskal-Wallis test followed by Dunn’s multiple comparisons test for non-parametric data.

### RNA isolation, library preparation, and sequencing

Third instar larvae of control (*elav>GFP RNAi*) and *deaf1* and *clamp* knockdown (*elav>deaf1 RNAi, elav>clamp RNAi*) genotypes were collected and their brains (optic lobes and VNC) were dissected in ice-cold PBS in rapid succession. Larval brains (n=12) were then immediately placed on ice in RNase-free 1.5ml tubes containing 100µl of TRIzol lysis reagent (Invitrogen, USA). We generated eight biological replicates for each genotype, with 12 brains per replicate. Larval brains were homogenized with a pestle. QIAGEN RNeasy Plus Universal Mini Kit was used for RNA extraction followed by the ZYMO RNA Clean & Concentrator-25 kit for further purification. RNA was eluted and confirmed for purity and concentration via 260/280 and 260/230 ratios measured using a Thermo Scientific NanoDrop One^C^. Illumina libraries were prepared and sequenced at Novogene Corporation for 150bp Poly-A selected mRNA and paired-end sequenced using an Illumina NovaSeq 6000, targeting at least 20 million reads per sample.

### RNA-seq analysis

Raw sequencing data were run for quality check using FastQC/0.11.9. Adapter content was trimmed with Trim-galore/0.6.10 and Cutadapt/4.6(*89*). Hisat2/2.2.1(*90*) was used for paired-end read alignment to the *Drosophila* reference genome dmel-r6.42, and Samtools/1.16.1(*91*) flagstat and idxstats for alignment checks. From Subread/2.0.2, featureCounts(*92*) was used to count reads at gene features with the following parameters: -p --countReadPairs -t gene -g gene_id -C -T 8. DESeq2/1.40.2(*93*) was used for differential gene expression analysis and log fold change shrinkage with ashr(*94*). We consider genes with padj (FDR) values below 0.05 to be differentially expressed. GO analysis was performed with gProfiler2/0.2.2(*72*) with the following parameters: organism = “dmelanogaster”, ordered_query = T, user_threshold = 0.05, correction_method = “g_SCS”. GO terms were grouped with rrvgo/1.12.2(*73*) by hierarchical parent term with threshold=0.96 and trimmed to only show biological process terms with uniqueness scores > 0.9. Odds ratio and Fisher’s exact test significance scores were calculated with the R package GeneOverlap/1.36.0(*85*).

### Phenotypic analysis of synapse number

Wandering third instar larvae were dissected in ice-cold Ca^2+^-free saline and fixed for 6 minutes in Bouin’s Fixative (Electron Microscopy Sciences, Cat# 15990). Dissections were washed several times with 1X PBS and then permeabilized with 1X PBS containing 0.1% Triton-X (Fisher Scientific, Cat# BP151-100). Dissected larvae were then blocked either for 1 hour at room temperature or overnight at 4°C in 1X PBS containing 0.1% Triton-X and 1% BSA (Sigma-Aldrich, Cat# A7906), followed by incubation in primary antibodies overnight at 4°C and secondary antibodies for 2-4h at room temperature or overnight at 4°C. Dissected larvae were mounted in Vectashield (Vector Laboratories, Cat# H-1000-10). The following antibodies were used at the indicated concentrations: mouse anti-Bruchpilot (Brp) at 1:100 (DSHB, Cat# nc82, RRID: AB_2314866), anti-Horseradish Peroxidase (HRP) conjugated to Alexa Fluor 647 at 1:500 (Jackson ImmunoResearch Labs, Cat# 123-605-021, RRID: AB_2338967), and Alexa Fluor 488 goat anti-mouse at 1:500 (Invitrogen, Cat# A-11029, RRID: AB_2534088). NMJs were imaged on a Nikon A1R HD confocal microscope with a 40x oil immersion objective. Boutons were quantified at the highly stereotyped larval NMJ4 in segments A2-A4.

Quantification of boutons was performed using the “multi-point” tool in FIJI/ImageJ (NIH, RRID: SCR_003070). To quantify individual active zones, we used Nikon Elements GA3 software to preprocess images with Guassian and Rolling ball filters and identify Brp puncta using the Bright Spots module. ROIs were drawn using HRP. All quantifications were performed masked to genotype. Violin plots were made with the R package ggplot2(*95*) and ggstatsplot(*96*). P-values were calculated with Welch’s t-test for unequal variances for single comparisons of parametric data or Kruskal-Wallis test followed by Dunn’s test and Benjamini-Hochberg p-value adjustment for multiple pairwise comparisons of nonparametric data.

### CUT&RUN

To identify genomic regions bound by DEAF1 and CLAMP in the larval brain, we performed Cleavage Under Targets and Release Using Nuclease (CUT&RUN(*53*, *54*)) on dissected third-instar *Drosophila melanogaster* larval brains. Third instar larvae of control (*elav>GFP RNAi*) and *clamp* and *deaf1* knockdown (*elav>clamp RNAi,elav>deaf1 RNAi*) genotypes were sexed for females prior to dissection and brains (optic lobes and VNC) were isolated in ice-cold PBS. For each biological replicate, brains from approximately 50 larvae were dissected in rapid succession and immediately transferred to 1.5 mL tubes containing 100 µL PBS on ice. Isolated larval brains were subjected to nuclei isolation prior to CUT&RUN. Briefly, tissue was gently homogenized and nuclei were released and purified using detergent-based lysis conditions optimized for neural tissue. Purified nuclei were then washed and resuspended in CUT&RUN Wash Buffer, and nuclei counts and viability were assessed using a Countess Automated Cell Counter. Each CUT&RUN sample contained >500,000 nuclei. Nuclei were then gently pelted and incubated with activated beads and resuspended in 100 µL of cold Antibody Buffer. CUT&RUN was performed using the CUTANA™ CUT&RUN Kit, Version 3 (EpiCypher, 14-1048) following the manufacturer’s protocol with minor modifications optimized for larval brain nuclei. Samples were incubated with antibodies against DEAF1 or CLAMP and IgG controls, and incubated overnight at 4 °C under gentle end-over-end rotation. All experimental samples represented independent biological replicates. Following MNase digestion and release of antibody-bound chromatin fragments, CUT&RUN DNA was purified for library preparation with the CUTANA™ Library Prep Kit (EpiCypher, 14-1001) according to the manufacturer’s instructions. Libraries were sequenced on an Illumina NovaSeq 6000 platform, generating 100 bp paired-end reads. On average, approximately 20 million reads per sample were obtained.

### CUT&RUN data analysis

Raw reads were processed with Trimgalore (0.6.10) to remove low-quality reads and clip adaptor sequences. Trimmed reads were aligned to the *Drosophila melanogaster* genome (Ensembl release 103, dm6) using Bowtie2 (v2.5.4)(*97*) with the parameters -x 1000 -- very-sensitive. picard/3.3.0 was used to remove any duplicates. Resulting BAM files were sorted and indexed with Samtools/1.16.192(*91*). Peaks were called with MACS3(*78*). For visualization, deepTools(*76*) bamCoverage was used to generate bigWig files for genome browser inspection and bedGraph files for peak calling. Regions of DEAF1 and CLAMP binding were identified using computematrix with normalization enabled. DEAF1 and CLAMP-bound regions were defined as those showing enrichment in the antibody samples, yielding 1,589 (CLAMP) and 6,721 (DEAF1) regions, which were annotated to genes using the ChIPseeker R package (v1.26.2)(*98*).

### Generation of average signal profiles from deepTools matrices

#### Generation of deepTools signal matrices

Genome-wide signal profiles were generated from CPM-normalized bigWig files. Signal matrices were computed using the deepTools computeMatrix function in reference-point mode. Profiles were centered on transcription start sites (TSSs) or synaptic DARs, with upstream and downstream distances set to span a total window of 2 kb. Signal was summarized in fixed-width bins of 100 bp), consistent with the resolution of the input bigWig files. Each bigWig file was processed independently against a set of predefined genomic regions corresponding to synaptic, early neuronal developmental and random nervous system gene sets. Synaptic,early neuronal developmental and random nervous system gene regions were provided as standard BED files containing chromosome, start, and end coordinates for each transcript. BED files were generated using the UCSC Table Browser. For each input, compressed matrix files (.mat.gz) were produced for downstream analysis.

#### Processing of matrix files and generation of average profiles

The resulting deepTools matrix files (.mat.gz) were processed in R for downstream analysis. For each genotype and gene-set group, replicate profiles were merged by averaging signal across replicates at the gene level for each bin. Gene-level profiles were then aggregated to generate summary profiles by calculating the mean signal across all genes at each bin position. Within each comparison, synaptic, early neuronal developmental and random nervous system regions were plotted separately. Mean signal was displayed as a line, with shaded regions representing ± standard error. The x-axis corresponds to position relative to the center of accessible regions, derived from bin indices and centered at the midpoint of the window.

### CUT&RUN and RNA-seq overlap with enrichment analysis

#### Gene universe and gene sets

A gene universe was defined from the *Drosophila melanogaster* dm6 genome using gene annotations extracted from the UCSC Genome Browser. Synaptic and random nervous system gene sets were used as defined earlier. Gene identifiers from these datasets were mapped to RefSeq mRNA identifiers using the org.Dm.eg.db annotation database via AnnotationDbi::select. For the CUT&RUN and RNA-seq overlap analysis, DEAF1-bound genes were identified based on CUT&RUN peak overlap with TSS-centered regions (±2 kb). For each condition, gene lists from two replicates were intersected to retain genes present in both replicates.

#### Overlap and enrichment analysis

Gene set overlap and enrichment were computed using the GeneOverlap R package(*85*). A GeneOverlapMatrix was constructed using the synaptic and control gene sets and the condition-specific gene lists, with the gene universe size specified as the background. Statistical significance of overlap was assessed using Fisher’s exact test. Effect sizes were quantified as odds ratios. P-values obtained from the overlap analysis were extracted and adjusted using the Bonferonni method. Adjusted p-values were used to assign significance levels. Odds ratios were extracted and plotted using ggplot2. Statistical significance was annotated using adjusted p-value thresholds (***<0.001; **<0.01; *<0.05; ns, not significant).

## Resource availability

### Materials availability

All unique/stable reagents generated in this study are available from the lead contacts.

### Data and code availability

Transcriptomic data have been deposited in the NCBI Gene Expression Omnibus (GEO) Database under the accession number GSE280405.

This paper also analyzes existing, publicly available data, accessible at doi: 10.1038/nature09715, doi:10.1126/science.abn5800, doi: 10.1016/j.devcel.2020.10.009, NCBI GEO Database GSE135810, GSE156455, GSE214707, GSE263159, GSE226514, and http://encodeproject.org.

Any additional information required to reanalyze the data reported in this paper is available from the lead contacts upon request.

## Acknowledgements

We thank the Developmental Studies Hybridoma Bank and Bloomington Drosophila Stock Center (NIH P40OD018537) for antibodies and fly stocks. FlyBase(*42*) provided access to critical reagent and genomic information and databases. We are grateful to Scott Gratz and Karla Kaun for invaluable assistance with figures and members of the Larschan and O’Connor-Giles labs, Oliver Hobert, and Melissa Harrison for thoughtful discussions and comments on the manuscript. This work was supported by a grant to KOG and EL from the National Institute of Neurological Disorders and Stroke, National Institutes of Health (R21NS125864) and a Brown University Carney Institute for Brain Science Zimmerman Innovation in Brain Science Award. James Kentro was supported by the NIH training grant T32GM136566 to the Brown Molecular Biology, Cell Biology, and Biochemistry Graduate Program, Gunjan Singh by Glenn Foundation for Medical Research Postdoctoral Fellowship, Glenn Foundation for Medical Research continuation Postdoctoral Fellowship and Audrey Medeiros by an NIH Predoctoral Individual National Research Service Award (F31NS122424).

## Declaration of interests

The authors declare no competing interests.

## Notes

### Competing Interest Statement

The authors have declared no competing interest.

### Summary of Updates

This version of the manuscript has been revised to include new CUT&RUN data providing mechanistic insight into DEAF1 and CLAMP regulation of synaptic genes, including direct DEAF1 binding and CLAMP-dependent DEAF1 occupancy at synaptic regulatory regions.

