## Supplemental_Figures for "Deaf1 regulates the coordinated repression of synaptic genes to limit synaptogenesis"

*Figure S1*

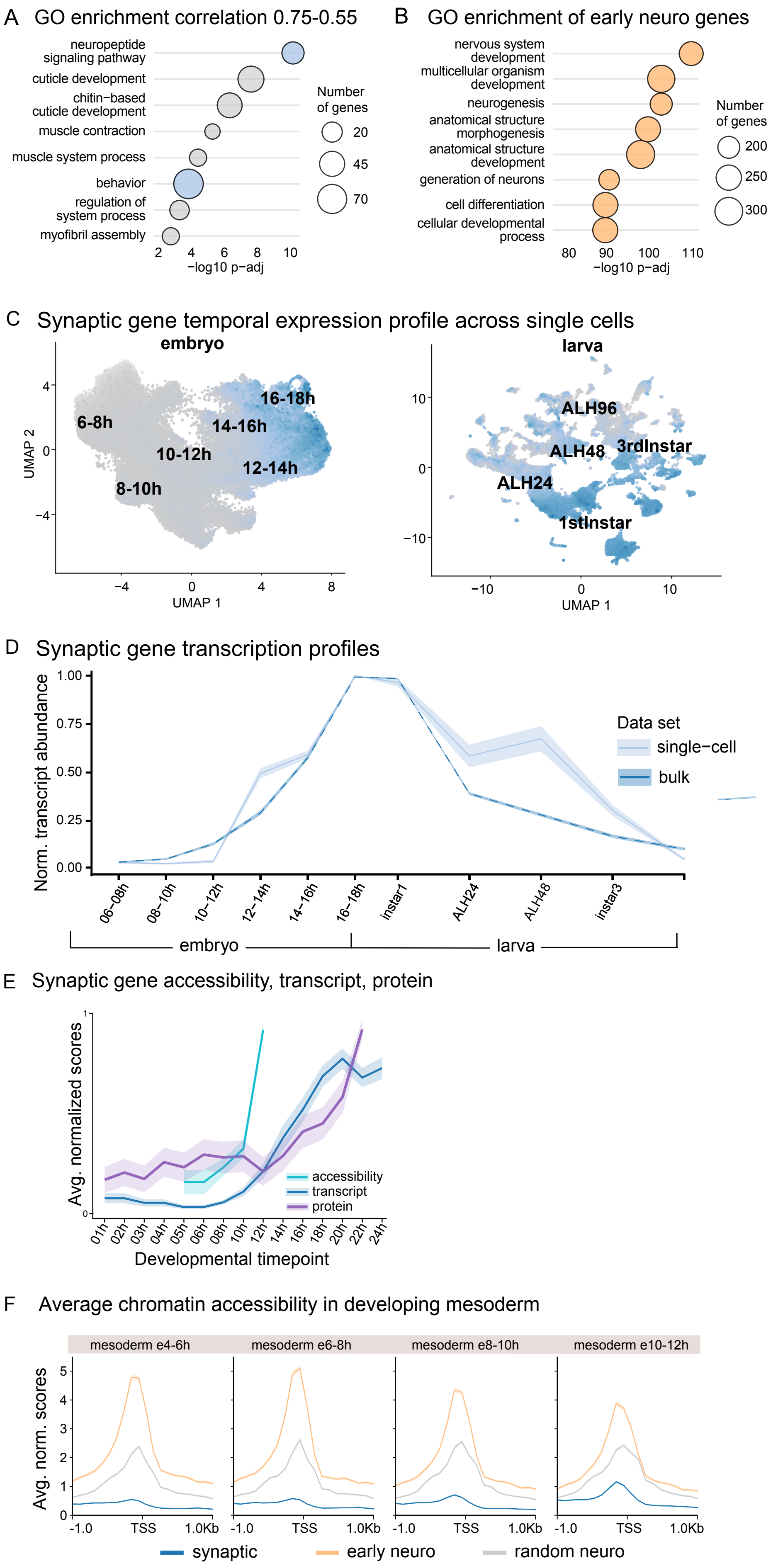
A GO enrichment correlation 0.75-0.55
neuropeptide signaling pathway
cuticle development
chitin−based cuticle development
muscle contraction
muscle system process
behavior
regulation of system process
myofibril assembly
Number of genes
20
45
70
2
4
6
8
10
−log10 p−adj
B GO enrichment of early neuro genes
nervous system development
multicellular organism development
neurogenesis
anatomical structure morphogenesis
anatomical structure development
generation of neurons
cell differentiation
cellular developmental process
Number of genes
200
250
300
80
90
100
110
−log10 p−adj
C Synaptic gene temporal expression profile across single cells
embryo
6-8h
8-10h
10-12h
12-14h
14-16h
16-18h
UMAP 2
UMAP 1
larva
ALH96
ALH48
3rdInstar
ALH24
1stInstar
UMAP 1
D Synaptic gene transcription profiles
Norm. transcript abundance
0.00
0.25
0.50
0.75
1.00
06−08h
08−10h
10−12h
12−14h
14−16h
16−18h
instar1
ALH24
ALH48
instar3
embryo
larva
Data set
single−cell
bulk
E Synaptic gene accessibility, transcript, protein
Avg. normalized scores
0
1
accessibility
transcript
protein
01h
02h
03h
04h
05h
06h
08h
10h
12h
14h
16h
18h
20h
22h
24h
Developmental timepoint
F Average chromatin accessibility in developing mesoderm
mesoderm e4-6h
mesoderm e6-8h
mesoderm e8-10h
mesoderm e10-12h
Avg. norm. scores
0
1
2
3
4
5
-1.0
TSS
1.0Kb
synaptic
early neuro
random neuro

*Figure S2*

A Enriched motifs in DARs

| Motif | ∑-log(*p*) | Best match |
|---|---|---|
| ACACAAAACA | 484 | pan |
| GGGGGGGTGG | 253 | l(3)neo38 |
| TCGCTTTTTT | 226 | sqz |
| GTTAAAAT | 186 | CG11617 |
| GAAATGAAAC | 180 | tll |
| GTGTTGCC | 168 | CG4854 |
| AGAAATTG | 149 | cad |
| TATATATATA | 141 | Cf2 |
| GGATGGGG | 125 | dar1 |
| CCAGTCAG | 118 | hth |
| CATGGCCA | 110 | pho |
| CGAAAGCG | 104 | Blimp1 |
| CCGCAGTATG | 103 | run |
| AGAGAGAG | 102 | GAF/Clamp |
| TGCAAAAT | 99 | rib |
| TTTTTTTTTCG | 97 | br |
| GTTTCTGT | 89 | kni |
| GTACATGACA | 84 | Vis |
| TTGCCAGTTT | 74 | prd |
| AGTGTGACCA | 45 | M1BP |
| GATTGTAGCG | 37 | Aef1 |
| AGTCCTGAGT | 35 | z |
| TCATTCAT | 32 | Dfd |
| TGGCACGTGC | 30 | $E_{spl}$mγ-HLH |
| GTGTATGTGTGT | 22 | cg |

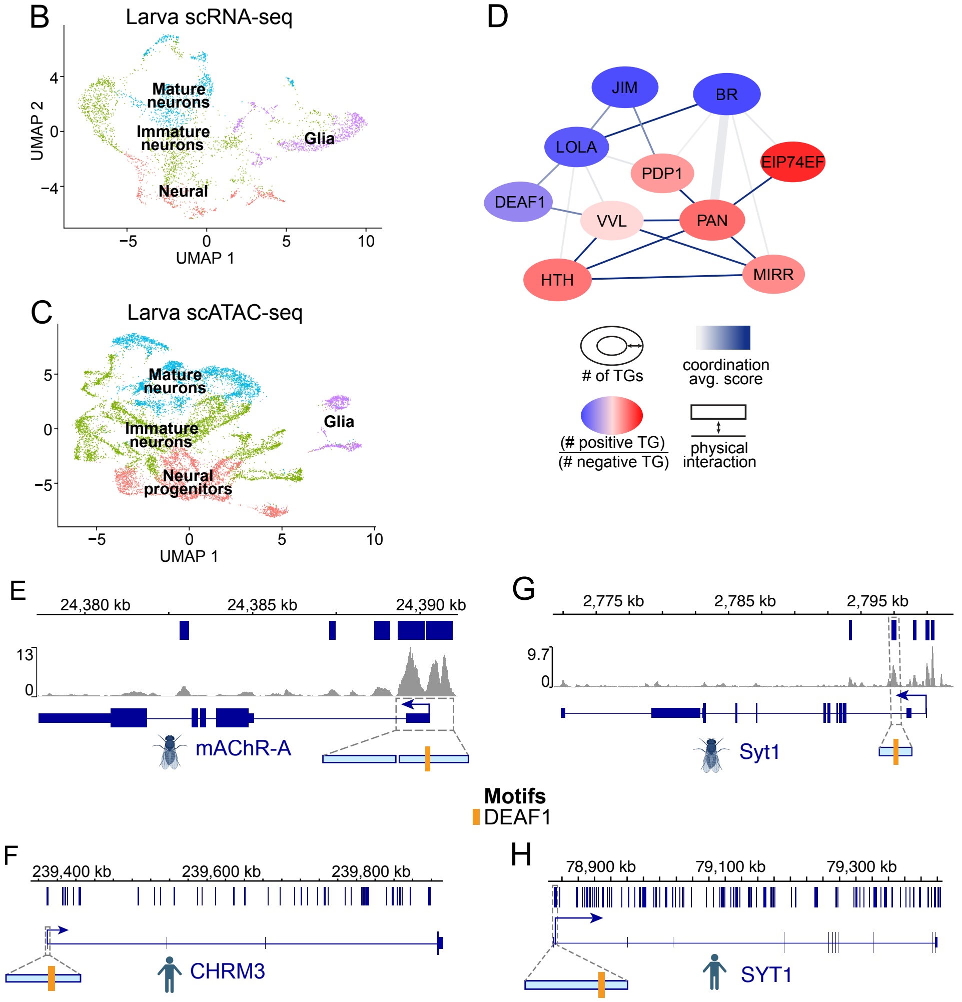

*Figure S3*

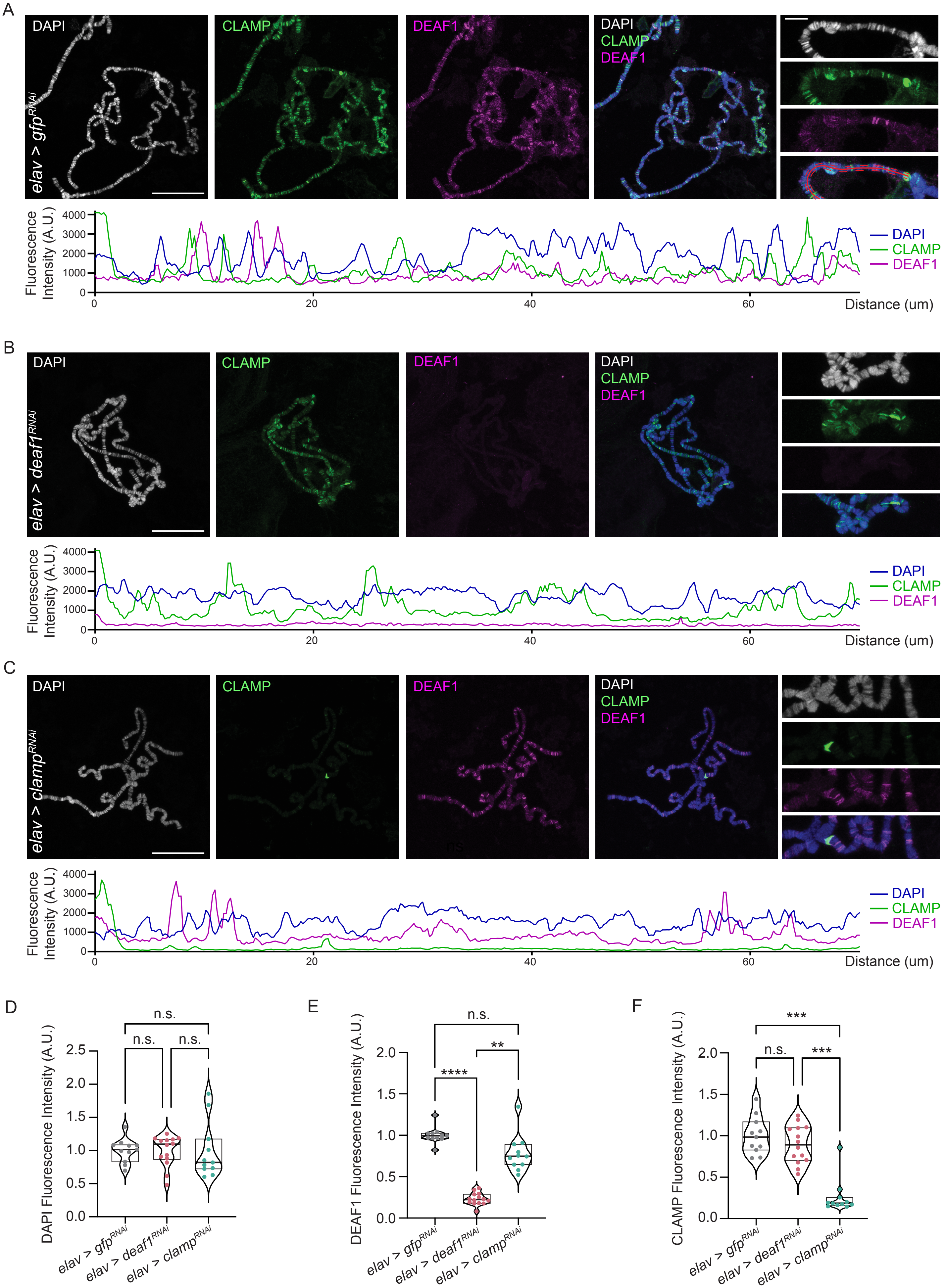
A
DAPI
CLAMP
DEAF1
elav > gfp^RNAi
Fluorescence Intensity (A.U.)
4000
3000
2000
1000
0
0
20
40
60
Distance (um)
B
elav > deaf1^RNAi
C
elav > clamp^RNAi
D
n.s.
n.s.
n.s.
DAPI Fluorescence Intensity (A.U.)
2.5
2.0
1.5
1.0
0.5
0.0
elav > gfp^RNAi
elav > deaf1^RNAi
elav > clamp^RNAi
E
n.s.
****
**
DEAF1 Fluorescence Intensity (A.U.)
F
***
n.s.
***
CLAMP Fluorescence Intensity (A.U.)

*Figure S4*

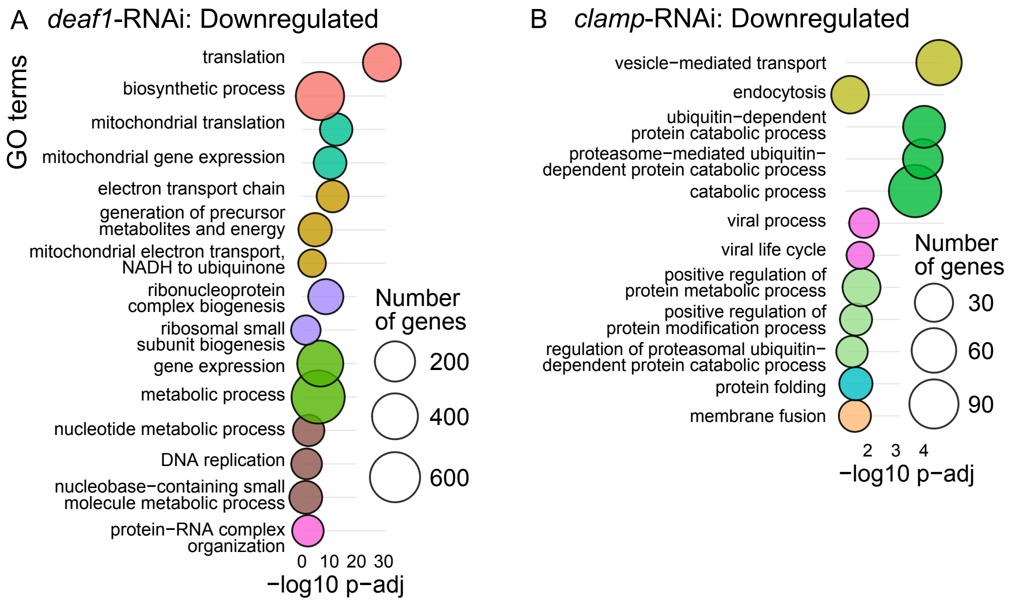
A deaf1-RNAi: Downregulated
GO terms
translation
biosynthetic process
mitochondrial translation
mitochondrial gene expression
electron transport chain
generation of precursor metabolites and energy
mitochondrial electron transport, NADH to ubiquinone
ribonucleoprotein complex biogenesis
ribosomal small subunit biogenesis
gene expression
metabolic process
nucleotide metabolic process
DNA replication
nucleobase−containing small molecule metabolic process
protein−RNA complex organization
0 10 20 30
−log10 p−adj
Number of genes
200
400
600
B clamp-RNAi: Downregulated
vesicle−mediated transport
endocytosis
ubiquitin−dependent protein catabolic process
proteasome−mediated ubiquitin−dependent protein catabolic process
catabolic process
viral process
viral life cycle
positive regulation of protein metabolic process
positive regulation of protein modification process
regulation of proteasomal ubiquitin−dependent protein catabolic process
protein folding
membrane fusion
2 3 4
−log10 p−adj
Number of genes
30
60
90

*Figure S5*

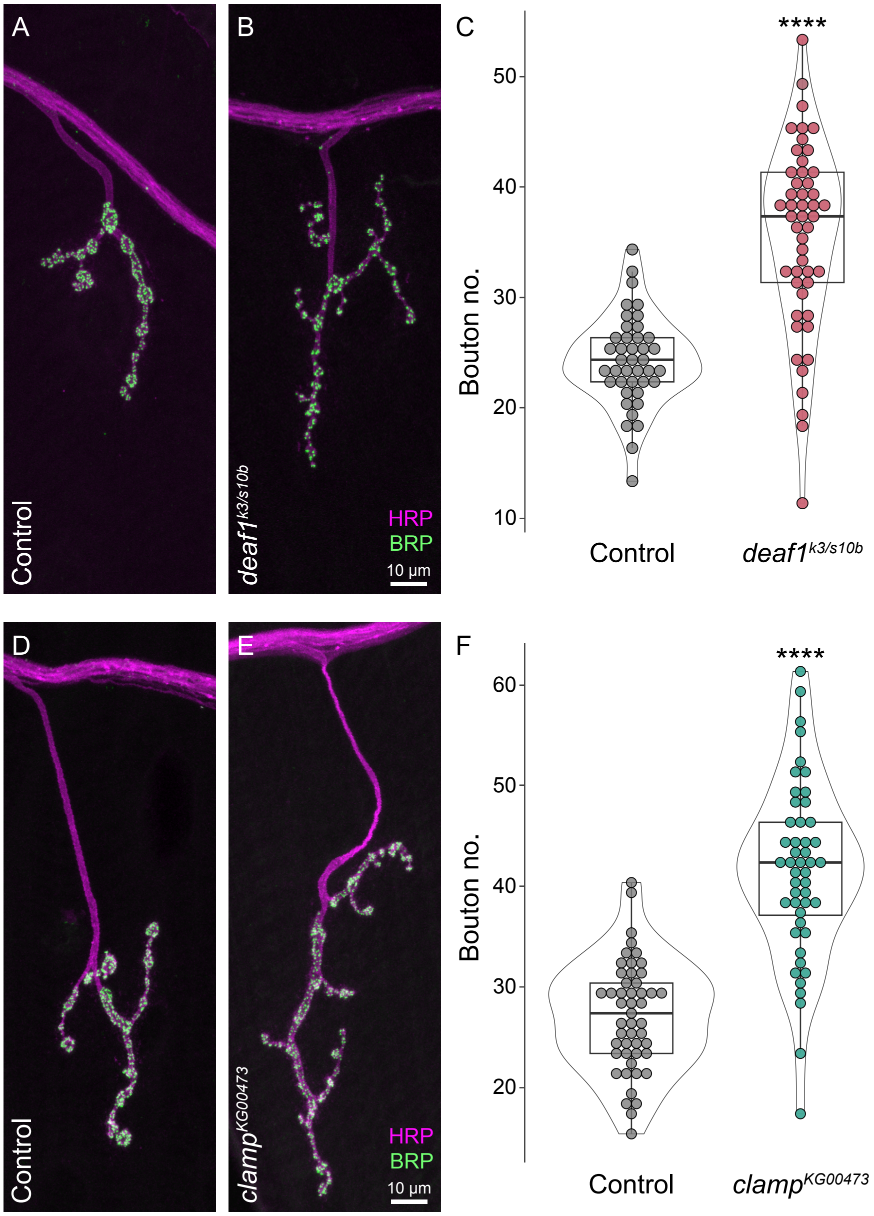

A
B
C
Control
deaf1k3/s10b
HRP
BRP
10 µm
****
Bouton no.
50
40
30
20
10
Control
deaf1k3/s10b
D
E
F
Control
clampKG00473
HRP
BRP
10 µm
****
Bouton no.
60
50
40
30
20
Control
clampKG00473

*Figure S6*

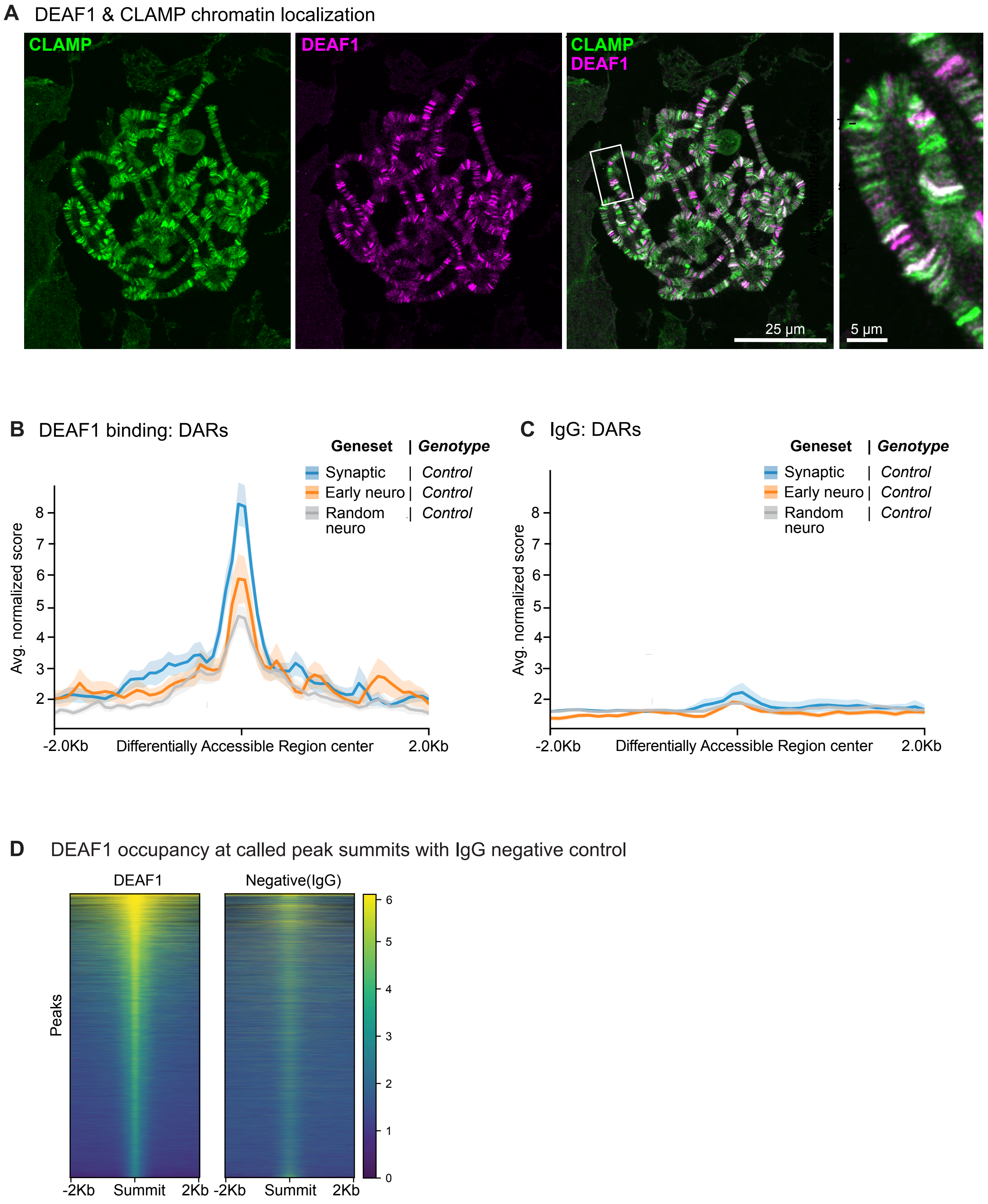

A DEAF1 & CLAMP chromatin localization
CLAMP
DEAF1
CLAMP
DEAF1
25 μm
5 μm
B DEAF1 binding: DARs
Geneset | Genotype
Synaptic | Control
Early neuro | Control
Random neuro | Control
Avg. normalized score
2
3
4
5
6
7
8
-2.0Kb
Differentially Accessible Region center
2.0Kb
C IgG: DARs
Geneset | Genotype
Synaptic | Control
Early neuro | Control
Random neuro | Control
Avg. normalized score
2
3
4
5
6
7
8
-2.0Kb
Differentially Accessible Region center
2.0Kb
D DEAF1 occupancy at called peak summits with IgG negative control
DEAF1
Negative(IgG)
Peaks
-2Kb
Summit
2Kb
-2Kb
Summit
2Kb
0
1
2
3
4
5
6

*Figure S7*

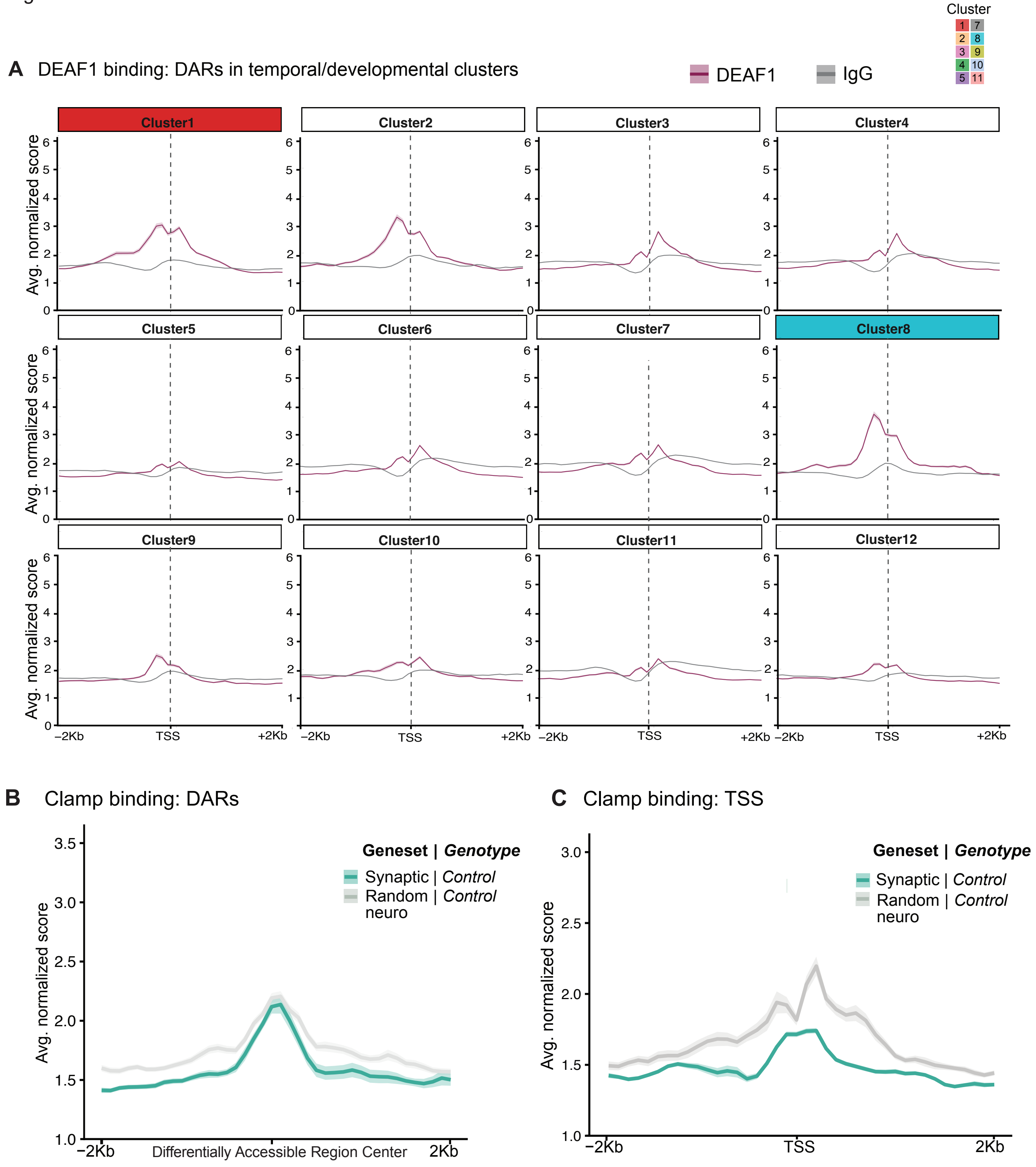
A DEAF1 binding: DARs in temporal/developmental clusters
DEAF1
IgG
Cluster
1 7
2 8
3 9
4 10
5 11
Cluster1
Cluster2
Cluster3
Cluster4
Cluster5
Cluster6
Cluster7
Cluster8
Cluster9
Cluster10
Cluster11
Cluster12
Avg. normalized score
−2Kb
TSS
+2Kb
B Clamp binding: DARs
Geneset | *Genotype*
Synaptic | *Control*
Random | *Control*
neuro
Avg. normalized score
−2Kb
Differentially Accessible Region Center
2Kb
C Clamp binding: TSS
Geneset | *Genotype*
Synaptic | *Control*
Random | *Control*
neuro
Avg. normalized score
−2Kb
TSS
2Kb
